# Large-scale volumetric two-photon calcium imaging enables cellular-resolution mesoscopic mapping of marmoset cortical areas

**DOI:** 10.64898/2026.05.28.728595

**Authors:** Santiago Otero-Coronel, David Grant Colburn Hildebrand, Tobias Nöbauer, Jeffrey Demas, Brandon Chen, Siegfried Weisenburger, Frank Tejera, Sverre Grødem, Kristian Kinden Lensjø, Guro Helén Vatne, Marianne Fyhn, Winrich A Freiwald, Alipasha Vaziri

**Affiliations:** Laboratory of Neurotechnology and Biophysics, The Rockefeller University, New York, NY, 10065, USA; Laboratory of Neural Systems, The Rockefeller University, New York, NY, 10065, USA; Price Center for the Social Brain, The Rockefeller University, New York, NY, 10065, USA; The Kavli Neural Systems Institute, The Rockefeller University, New York, NY, 10065, USA; Center for Integrative Neuroplasticity, Department of Bioscience, University of Oslo, Oslo, Norway

**Keywords:** two-photon microscopy, large-scale imaging, mesoscopy, volumetric calcium imaging, temporal multiplexing, Light Beads Microscopy (LBM), ribosome tethered jGCaMP8 ribo-jGCaMP, common marmoset Callithrix jacchus, non-human primate, Middle Temporal area MT, direction tuning, auditory cortex

## Abstract

Primate neocortex is organized into functionally defined domains. Understanding how the functional organization within each domain emerges from single-cell tuning properties has been hampered by the lack of imaging tools that can capture large-scale activity at different depths of the scattering primate brain at cellular resolution and physiological timescales. Here, we demonstrate mesoscopic and volumetric cellular-resolution calcium imaging of entire cortical areas in the marmoset brain at multi-Hertz rates by combining fluorescence lifetime-limited temporally multiplexed two-photon recording based on Light Beads Microscopy (LBM) with optimized calcium indicator targeting. Using this approach, we perform volumetric recording of neuronal population activity within cortical volumes spanning up to 4 × 4 × 0.3 mm^3^ capturing the simultaneous activity of over 96,000 cells, including more than 46,000 stimulus-responsive neurons. In auditory cortex, we found fine-grain heterogeneous tuning of individual neurons to frequencies within broader tonotopic gradients. In the middle temporal area (MT) we found direction tuning along functional continua rather than discrete categories, encompassing uni-, bi-, and pandirectional tuning, where cells with similar tuning tend to cluster. Tuning direction follows a columnar organization with a lateral periodicity of ∼500 µm and low variability across depth. Columns are organized by a pinwheel structure. Noise correlations between cells, revealing functional interactions, decay isotropically and monotonically with distance but remain significant even at lateral distances exceeding 3 mm and thus the scale of multiple columns. Together, these discoveries reveal the need for neurotechnologies bridging scales from single cells to multi-columnar systems. The technological advancements open the door for future large-scale, cellular-resolution investigations of functional organization and population dynamics across entire cortical areas in the primate brain.

## INTRODUCTION

The history of systems neuroscience has been shaped by the attempt to uncover the functional organization of brain regions, from the tuning of single cells to the large-scale overall architecture^1–3^. Different levels of organization have been investigated with different techniques. Single cell properties, for example, have been studied with extracellular electrophysiological recordings, and functional maps have been obtained using imaging approaches^4–6^. Results from these different approaches could then be mapped onto each other^7,8^. With these approaches, the classical picture of the visual system has been stitched together. At the same time, the need for technology to bridge all levels of functional organization and connect this picture with an understanding of neural dynamics unfolding within it, has long been recognized ^9–11^, yet their availability and applicability in the large brains of non-human primates (NHP) have remained limited^12,13^. The key challenge is to develop the capability of simultaneously recording the activity of large neuronal populations across extended cortical territories while preserving single-cell resolution.

Recently, recordings from large neuronal populations at cellular resolution with high signal-to-noise ratio (SNR) and at physiological time-scales^11,14^ have become possible thanks to advances in optical neuro-recording technologies, including cellular-resolution optical mesoscopy platforms^10^, spatio-temporal multiplexing methods^11,15–23^ and other volumetric recording approaches^24,25^, coupled with the development of bright and sensitive calcium indicators^26^ and their targeted localization^27–29^. To date, due to their relatively smaller brain size, broad accessibility and a well-established genetic tool kit, these technical advances have been mainly implemented and optimized in mice. However, fundamental differences in cognitive, behavioral, and sensory capabilities as well as brain organization require direct investigations of the primate brain. Compared to primates, mice have lower visual acuity^30–32^, lack foveal vision^33,34^, and predominantly exhibit a ‘salt-and-pepper’ rather than columnar^35^ organization in visual areas (see ref. ^36^ cf. ^37,38^). They also lack key primate specializations including invariant object recognition^39^, higher visual areas that are specialized for motion^40^ and face^41^ processing, and observational learning through imitation^42^. NHP models therefore remain essential for studying a broad range of questions in systems neuroscience.

In NHPs, neuronal tuning and function vary across brain lobes, cortical areas, columns, and neighboring cells. Therefore, given the relatively large size of NHP brains^43^, the spatial location and tuning of neurons are intertwined at scales that can span from centimeters to micrometers. As a result, this relationship has been classically studied using methods that provide either large-scale spatial coverage, such as intrinsic signal optical imaging (‘intrinsic imaging’) and functional magnetic resonance imaging (fMRI), or single-cell resolution, such as electrophysiology. However, a technique that simultaneously offers high-throughput, single-cell resolution recording of neuronal population activity across entire primate cortical areas remains to be implemented. Mesoscopic two-photon (2p) calcium imaging is uniquely positioned to bridge this gap by enabling direct visualization of neuronal activity at single-cell resolution across extended cortical territories.

At the same time, calcium imaging in NHPs presents unique challenges due to their thicker skulls, larger brains, cortical folding, higher cortical thickness and light scattering, and lack of general availability of transgenic lines expressing calcium indicators^44,45^. In addition, primate cortical layer I is largely devoid of somata^46,47^ and is thicker compared to mice. Thus, obtaining access to neurons in cortical layers II/III, where typically most recordings are performed, is challenging. Other limiting factors include reduced viral transduction efficiency^48,49^, constraints on optical access imposed by chronic recording requirements^50^, and the specialized infrastructure inherent to long-term NHP experimentation.

On this background, the marmoset monkey presents a promising NHP model, offering some of the technical advantages of rodent models in a species where the anatomy, functional organization, cognition, and behavioral repertoire of primates can be studied. Marmosets can be trained to fixate^34^, perform object recognition tasks^39^, and they exhibit rich social behaviors similar to those of larger primates^51^. Notably, the 7.5 cm^3^ volume of the marmoset brain is about one order of magnitude smaller than that of larger primates such as macaques^52^, while exhibiting a relatively flat cortical surface that offers higher optical accessibility^51^. Leveraging this trait, calcium imaging studies have been performed in the marmoset somatosensory^53–57^, visual^12^, auditory^58–60^, and motor^61^ cortices. Further, tetracycline-regulated promoter systems that amplify and provide temporal control over gene expression have been established and leveraged for long-term calcium imaging^53^. Nonetheless, the state of the art in marmoset calcium imaging is single-plane recordings over submillimeter fields of view (FOVs) with limited SNR resulting in tens to low-hundreds of recorded cells ^12,53^. This stands in sharp contrast to the to-date largest volumetric calcium recordings in mice spanning more than a 15 cubic millimeter volume with more than 1 million active neurons^11,14^.

Here, we present an optimized and scalable volumetric imaging platform that integrates advancements in large-scale cellular resolution neuronal population calcium imaging with the latest calcium indicators and their targeted expression in the cortex of marmoset monkeys. Designing viral vectors optimized for delivery of improved jGCaMP8 constructs to primate neurons and strong, soma-targeted expression with reduced neuropil contamination and background fluorescence facilitated improved single-cell detection in marmosets. We combined these molecular improvements with Light Beads Microscopy^11^ (LBM), a highly optimized volumetric 2p imaging platform that allows for fluorescence-lifetime-limited pixel acquisition rates of up to ∼150 MHz on a mesosocopic scale.

We performed volumetric neuronal population recordings within marmoset visual and auditory cortex spanning volumes up to 4 × 4 × 0.3 mm^3^ at single-cell resolution, simultaneously capturing the activity of over 46,000 stimulus-responsive cells at 1.9 Hz. To our knowledge, our work represents the first demonstration of volumetric imaging of neuronal population activity in a NHP. By recording cellular-resolution activity across cortical volumes of up to ∼5 mm³, we extend the previously reported number of simultaneously recorded neurons in NHPs^12,62^ by more than two orders of magnitude.

Leveraging our platform, we explored mesoscale functional organization across multiple cortical areas in the marmoset. In visual area MT, we observed a volumetric spatial organization of direction-tuned neurons in pinwheel motifs tiled laterally with a ∼500 µm spatial periodicity, a columnar organization across the recorded cortical depth, and long-range lateral noise correlations spanning multiple millimeters. Population-level analysis revealed that direction tuning properties in MT form a continuum, encompassing unidirectional, bidirectional, and pandirectional responses, rather than falling into discrete, separate classes. In auditory cortex, we found fine-grained heterogeneous tuning of individual neurons to frequencies within broader tonotopic gradients, revealing a more granular organization than previously appreciated. Recordings spanning the transition between visual and auditory areas uncovered neurons responsive to both modalities, indicating that functional boundaries can contain intermixed multimodal cells. Our results demonstrate the capabilities and the biological utility of our platform and establishes it as a powerful tool for functional dissection of the fine-scale and long-range principles of primate cortical organization at single-cell resolution and unprecedented scale.

## RESULTS

### Viral vector and indicator design for calcium imaging in marmosets

The expression level, brightness, kinetics, and localization of calcium indicators directly affect the quality of the recorded neuronal signals and potential for neurotoxicity. While many mouse transgenic lines expressing calcium indicators are commercially available, marmoset transgenic lines were only generated^63,64^ relatively recently and are not readily available to the broader community. As a result, calcium imaging experiments in marmosets have relied on injections of dyes^57–59^ or viruses^53,56,55,12^ to achieve indicator expression. However, viruses developed and optimized for mice cannot necessarily be used in NHPs. This is for a number of reasons. First, viral constructs with capsids well tolerated in mice can elicit stronger immune responses in NHPs, increasing the risk of local neurotoxicity and reducing transgene expression^65,66^. Second, constructs established to produce robust fluorescence levels in mice in vivo can produce expression levels that are either undetectable or cytotoxic in marmosets^53^. Third, constructs of sufficient brightness for imaging in mice, may not generate sufficiently strong signals in the primate brain with increased light absorption due to its elevated lipofuscin content^67^ and increased cortical thickness. Finally, large-scale and volumetric calcium imaging must be performed within safe biological limits of brain exposure to average laser power and pulse energy, further increasing the need for more sensitive and brighter sensors to obtain neuronal activity recordings with sufficient SNR. These challenges are compounded by the inherently resource-intensive nature of NHP experiments, which typically involve extended training and longitudinal recording paradigms. Consequently, the development and implementation of expression systems that deliver optimized calcium indicators providing strong, controllable, and non-toxic expression are essential for enabling large-scale, volumetric recordings in marmosets.

One technical advance for safeguarding neuronal overexpression of calcium indicators in marmosets is the tetracycline-responsive transactivator (Tet) system, which allows controllable amplification of transgene expression to levels compatible with in vivo calcium imaging and has been successfully used in the marmoset monkey^53^. The second advance consists in recent improvements in calcium indicator brightness, kinetics, and subcellular confinement, which have been widely adopted in rodent studies. Yet, they still require integration with primate-compatible capsids and promoters followed by empirical validation of their performance before they can be widely adapted and utilized by the NHP community. These improvements include the development of jGCaMP8^26^ sensors for NHPs, which has been shown in mice to exhibit increased brightness and faster response kinetics as well as soma-targeted^28^ and ribosome-tethered^29^ calcium indicator variants with expression profiles primarily restricted to cell bodies. Such localized expression of calcium indicators has been shown to reduce neuropil signal contamination leading to improved single-neuron detectability and higher neuronal signal-to-background ratios, although at the cost of reduced overall brightness^29^.

To compare the trade-offs and evaluate the suitability of these calcium indicators for large-scale imaging in marmosets, we designed viruses to deliver cytosolic jGCaMP8s (cyto-jGCaMP8s), soma-targeted^28^ jGCaMP8s (soma-jGCaMP8s), and ribosome-tethered^29^ jGCaMP8s (ribo-jGCaMP8s) under Tet control and compared them using 2p mesoscopic imaging in the marmoset cortex. To this end, we performed through-skull intrinsic imaging^68^ and presented moving-versus-static dots visual stimulus contrast (STAR Methods) to localize motion-sensitive area MT. We then implanted a 10 mm-diameter cranial window chamber covering a large lateral region of the temporal lobe containing area MT. Within the window, we identified three cortical regions that were separated by large blood vessels, into which we injected cyto-jGCaMP8s in 15 injection sites, soma-jGCaMP8s in 16 injection sites, and ribo-jGCaMP8s in 9 injection sites (**Figure 1A**).

**Figure 1.**
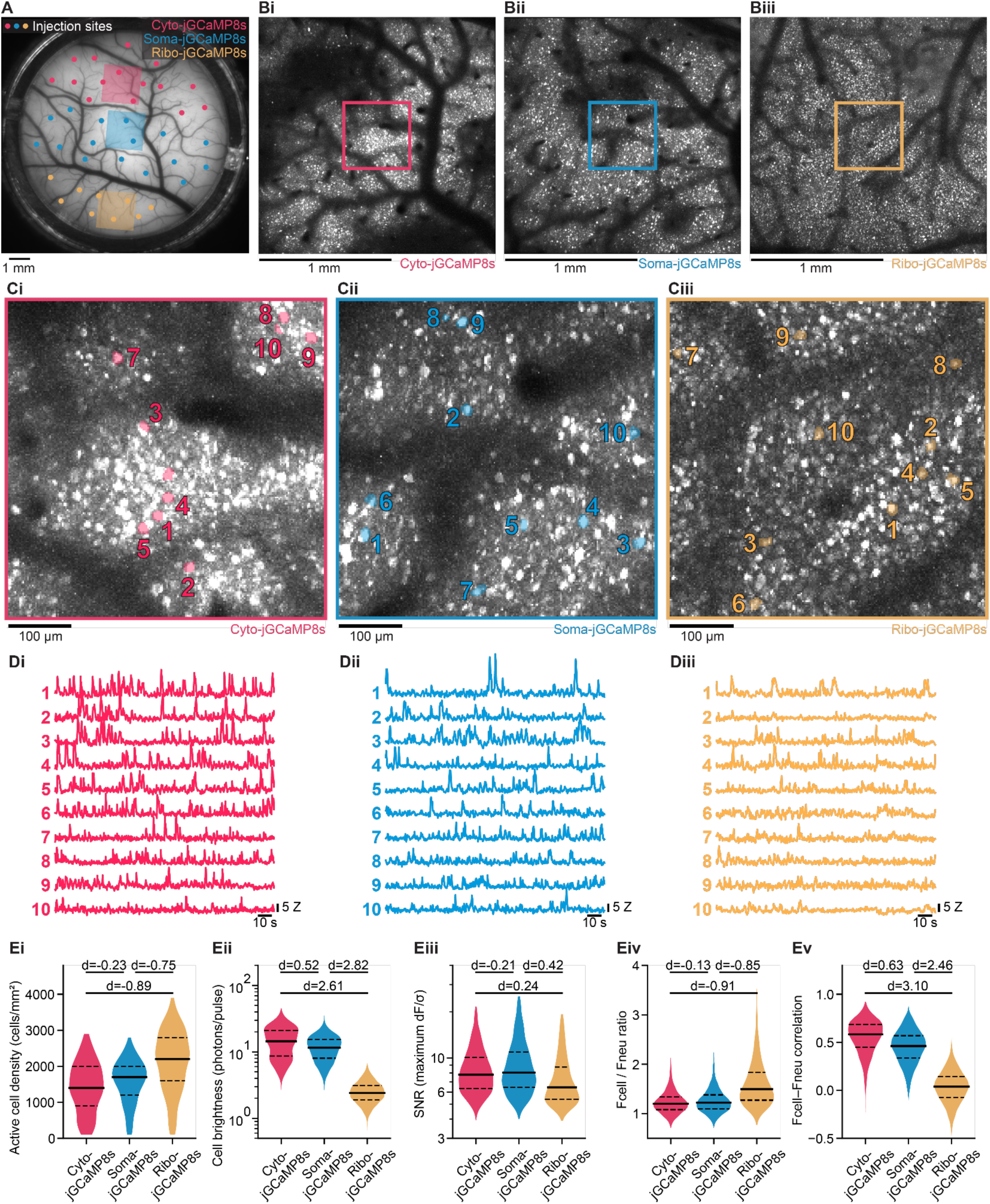
Cellular-resolution 2p mesoscopic imaging and comparison of cyto-, soma-, and ribo-jGCaMP8s in marmosets. A. Optical window (10 mm diameter) implanted over visual area MT. Small circles indicate viral injection sites for cyto-jGCaMP8s (red), soma-jGCaMP8s (blue), and ribo-jGCaMP8s (yellow). Shaded squares denote the size-matched 1.8 × 1.8 mm² regions shown in B and used for quantitative comparisons in E. B. Mean images of 1.8 × 1.8 mm² regions expressing (i) cyto-jGCaMP8s, (ii) soma-jGCaMP8s, and (iii) ribo-jGCaMP8s, illustrating differences in labeling density and neuropil signal. Colored rectangles indicate the 500 × 500 µm² subregions enlarged in C. C. Enlarged 500 × 500 µm² subregions from B for (i) cyto-jGCaMP8s, (ii) soma-jGCaMP8s, and (iii) ribo-jGCaMP8s, with footprints of ten representative cells highlighted for each indicator. D. Calcium activity traces of the ten cells highlighted in C, showing high-SNR transients for (i) cyto-jGCaMP8s, (ii) soma-jGCaMP8s, and (iii) ribo-jGCaMP8s. E. Quantitative comparison of the three indicators across cells detected in the regions shown in B, including (i) active cell density across 100 × 100 µm² patches, (ii) cell brightness, (iii) SNR (maximum of the noise-normalized activity trace, dF/σ), (iv) ratio of mean soma to neuropil fluorescence (F_cell_ / F_neu_), and (v) soma-neuropil Pearson correlation (F_cell_ – F_neu_). All panels show violin plots with median (solid line) and first and third quartiles (dashed lines). Cohen’s d values indicate the effect size between indicators for each metric (small: |d| < 0.2; medium: 0.5 ≤ |d| ≤ 0.8; large: |d| > 0.8).

To monitor the expression levels of all three regions injected with the different viral constructs, we performed single-plane 2p imaging using a mesoscope^10^ that provided access to FOVs of up to 6 × 5.4 mm^2^ at an excitation numerical aperture (NA) of 0.6 and a collection NA of 1.0. We recorded from layer II/III of marmoset MT at 200 µm depth while capturing spontaneous activity and responses to visual stimuli consisting of white and black dots coherently moving over a grey background 16 different directions (dots stimulus, STAR Methods; **Figure S1**). On day 14 after viral injections, neurons exhibiting strong fluorescence accompanied by transient fluctuations were observed across the optical window (**Figure 1B–D, Videos S1–S3**). Regions injected with cyto-jGCaMP8s and soma-jGCaMP8s exhibited higher fluorescence levels and signal amplitudes than regions injected with ribo-jGCaMP8s, where dynamics were visible and expression was confined to the cell bodies.

To quantify these observations, we selected a size-matched 1.8 × 1.8 mm² area within each of the regions and imaged at a pixel spacing of 2.5 µm and frame rates ranging from 2.6 to 3.6 Hz using an average laser power of 30 mWWe used the suite2p package^69^ to perform non-rigid motion correction and automated anatomical cell segmentation, extracting raw fluorescence values (F_raw_) from individual cell regions of interest (ROIs) as well as from their surrounding neuropil-containing pixels (F_neu_) and obtain the neuropil-corrected cell fluorescence (F_cell_). We defined the activity signal as dF_cell_ = F_cell_ − F_0_, and the noise-normalized activity signal as Z = dF_cell_/σ_cell_, where σ_cell_ represents the noise estimation for each cell (STAR Methods). For all quantifications throughout this and subsequent experiments, segmented ROIs were classified as true, active cells if the 99^th^ percentile of their activity Z, and the 99th percentile of their dF_cell_/σ_rec_, were both greater than 3, where σ_rec_ denotes the recording-specific noise (STAR Methods). Across 1.8 × 1.8 mm² imaging areas, we detected 4,051 active cells in the region injected with cyto-jGCaMP8s (1,267 cells per mm²), 4,858 in the soma-jGCaMP8s region (1,519 cells per mm²), and 6,624 in the ribo-jGCaMP8s region (2,066 cells per mm²). As local cell densities can be affected by the presence of large blood vessels and injection sites, potentially introducing variability across analyzed regions, we repeated this analysis by subdividing each region into 100 × 100 µm² subregions and excluding subregions without any active cells, yielding similar results (**Figure 1Ei**).

We then assessed how the different construct designs influenced overall brightness and SNR. Cells in regions injected with cyto-jGCaMP8s exhibited the largest F_cell_ (15.32 ± 7.80 photons per pulse, mean across cells ± SD, STAR Methods), closely followed by soma-jGCaMP8s (12.00 ± 4.98 photons per pulse), whereas cells expressing ribo-jGCaMP8s (2.61 ± 1.00 photons per pulse) were ∼5-fold dimmer (**Figure 1Eii**). To see if the different brightness of the indicators resulted in different SNR levels, we quantified the SNR of each cell as the maximum value of the noise-normalized Z activity for each cell. We found SNR was highest for cells expressing soma-jGCaMP8s (9.74 ± 5.03), followed by cyto-jGCaMP8s (8.82 ± 3.55), and then ribo-jGCaMP8s (7.89 ± 3.99) (**Figure 1Eiii**). Thus, despite a 5-fold lower brightness, ribo-jGCaMP8s led only to a 20% lower SNR while yielding 30–60% more active cells than soma-jGCaMP8s and cyto-jGCaMP8s. Consequently, the lower SNR of the population of ribo-jGCaMP8s expressing cells may reflect the inclusion of cells whose low signal would be lost in the background and thus fail to be detected when using soma-jGCaMP8s or cyto-jGCaMP8s.

When analyzing the neuropil signals, ribo-jGCaMP8s expression was highly confined to the cell body, resulting in the highest F_cell_ / F_neu_ ratio (1.62 ± 0.50), followed by that of soma-jGCaMP8s (1.27 ± 0.25) and cyto-jGCaMP8s (1.23 ± 0.25) (**Figure 1Eiv**). In addition, the Pearson correlation between F_cell_ − F_neu_ signal was lowest and near zero for ribo-jGCaMP8s (0.03 ± 0.16), and much higher for soma-jGCaMP8s (0.45 ± 0.17) and cyto-jGCaMP8s (0.56 ± 0.18) indicating that the ribosome tethering reduced the contribution of neuropil-derived signals to the somatic signal (**Figure 1Ev**).

Overall, these results demonstrate for the first time in an NHP that mesoscopic imaging with efficient excitation and collection, combined with the brighter and faster jGCaMP8 calcium indicators, can capture the population dynamics of thousands of neurons with high SNR over a multi-millimeter FOV. Our above results are consistent with prior findings in mice^29^, where ribo-jGCaMP8s has been found to greatly reduce neuropil contamination and improve cell detectability though at the cost of lower fluorescence yield. Accordingly, achieving SNR levels with ribo-jGCaMP8s that are comparable to cyto-jGCaMP8s and soma-jGCaMP8s required ∼ 70-100 % more average power. This leads to a dual strategy: the reduced neuropil contamination and higher neuronal yield provided by ribo-jGCaMP8s can be exploited in single-plane recordings, where the above ∼2-fold higher power penalty remains within established safe power exposure limits ^70,21,22,11^. However, for volumetric recordings using LBM that need more total power, using ribo-jGCaMP8s may exceed safe biological exposure limits. Therefore, volumetric recordings using LBM in marmosets require the use of the brighter soma-jGCaMP8s and cyto-jGCaMP8s indicators.

### Cellular-resolution mesoscopic calcium imaging in marmoset visual area MT

Understanding how neuronal tuning is organized across large cortical territories requires imaging technologies that combine large, volumetric multi-millimeter FOVs recording capability across cortical depths with single-cell spatial resolution. Spanning these scales is particularly important for primate brains, which have relatively large cortical areas whose functional organization exhibits complex spatial patterns spanning micrometers to centimeters. The recent development of large-scale optical imaging platforms in rodents that bridge cellular and mesoscopic scales^10,71,23^ has enabled the functional characterization of thousands of individual cortical neurons across millimeter-scale. However, to our knowledge no previous study has described the spatial organization and functional tuning of simultaneously recorded neuronal populations over such large areas at the cellular scale in a primate.

Previous studies in primates using intrinsic imaging have shown that MT^72,73^ contains repeating columns tuned to movement directions, analogous to how columns are tuned to edge orientations in V1^5,74,75^. Intrinsic imaging in galago monkeys^76^ and electrophysiological recordings in capuchin monkeys^73^ suggest that in MT these columns are arranged around spatial centers to form pinwheel-like patterns with sharp linear discontinuities (fractures) in preferred direction. Classical electrophysiological experiments in macaques^77^ and recent 2p calcium imaging of inhibitory neurons in marmosets^12^ have both found cellular-resolution evidence consistent with such organization of direction tuning in area MT, where neurons tuned to the same motion direction tend to cluster together. However, the relationship of single cell tuning and the overall functional organization of area MT have not before, to our knowledge, been determined simultaneously with the same method and within the same subject.

To target our imaging windows to MT, we first coarsely localized it using through-skull intrinsic imaging while presenting awake, head-restrained marmosets with the same moving-versus-static dots visual stimuli used to localize MT for calcium indicator experiments. MT was identified as a region with stronger hemodynamic responses to the moving dots than to the static dots (STAR Methods), and the accuracy of the optical window implantation targeting was then confirmed using though-window intrinsic imaging with the same stimulus (**Figure 2A–B**; **Figure S3**). We then used our custom-built large-scale imaging platform (STAR Methods) to perform 2p mesoscale calcium imaging of a region laterally encompassing MT in animals expressing ribo-jGCaMP8s. We performed single plane cellular resolution recordings from layer II/III of MT at 200 µm depth within a FOV of 2.6 × 2.5 mm^2^ at 3.35 Hz, and confirmed dense, soma-confined expression of jGCaMP8s (**Figure 2C–D**). Individual neurons exhibited strong activation in response to motion stimuli, reliably reproduced across the ten repetitions of each motion direction (**Figure 2E**). Within this FOV, we detected 10,534 total active cells. Here and in all subsequent experiments, we classified stimulus-responsive cells as the subset of active cells for which an ANOVA indicated that Z activity values differed between motion and blank conditions (p < 0.01) and responded to at least one motion direction with an average Z activity larger than 1 (STAR Methods). Among the active cells, we found 4,492 (42.6%) stimulus-responsive cells.

**Figure 2.**
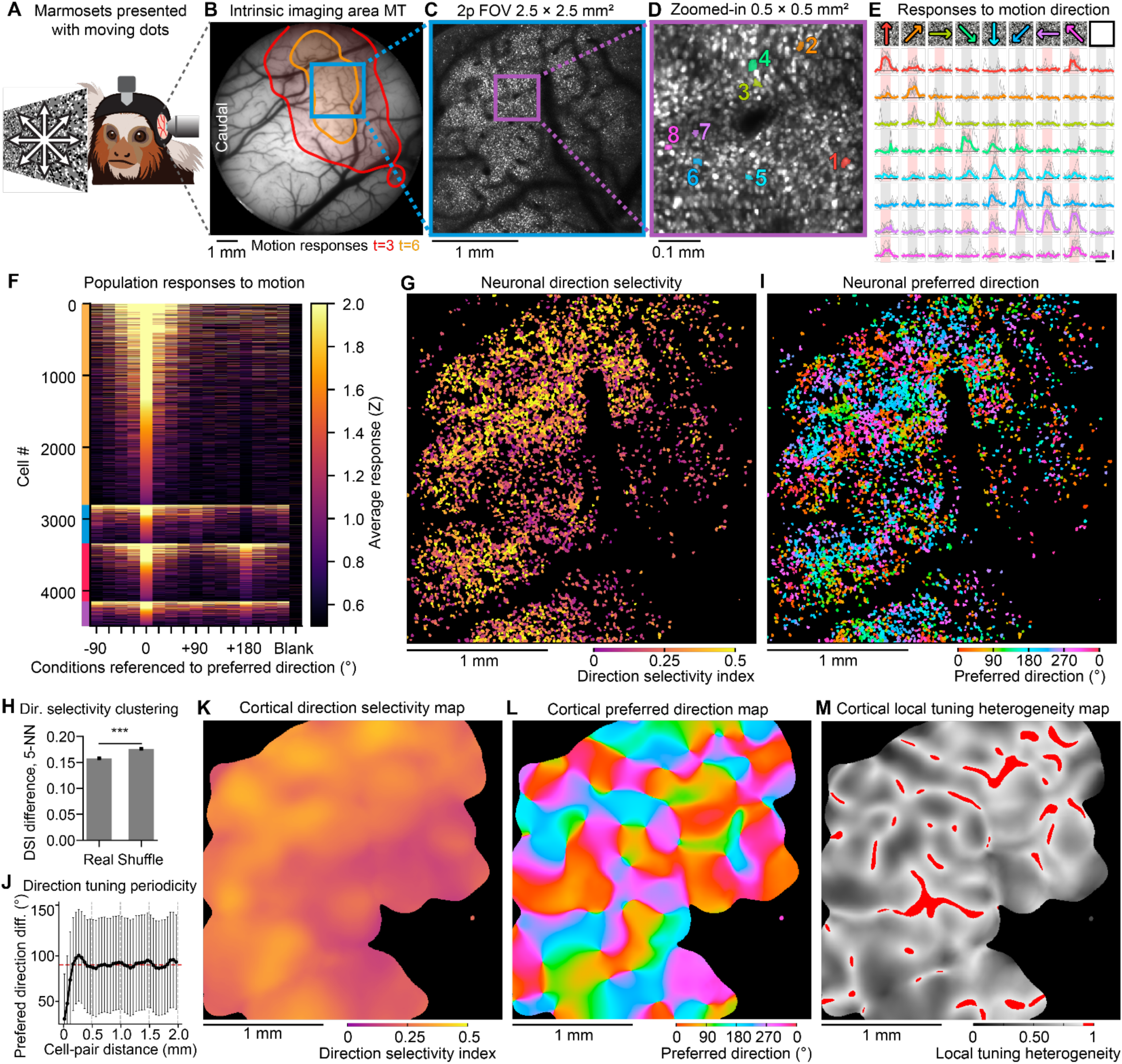
Cellular-resolution mesoscopic imaging and direction tuning in marmoset MT. A. Experimental setup. Head-restrained, awake marmosets were presented with moving-dot visual stimuli while intrinsic optical imaging or 2p calcium imaging was performed. B. Functional map overlaid on a brightfield image of the cranial window showing the location of MT based on responses in intrinsic imaging to a moving-dots stimulus. The blue rectangle marks the mesoscopic 2p FOV shown in panel C. C. Mean image of a mesoscopic 2p recording showing dense expression of ribo-jGCaMP8s throughout the entire FOV of 2.5 × 2.5 mm². The purple rectangle marks the zoomed-in region shown in panel D. D. Close-up view of a 0.5 × 0.5 mm² region from panel C shows dense labeling and soma-confined expression of ribo-jGCaMP8s. The footprints of 16 segmented cells are numbered and colored based on their preferred direction (hue-to-direction reference in 1E). E. Stimulus-aligned activity for the 16 cells shown in panel D displays stimulus-locked responses to different subsets of moving-dots directions. Each row corresponds to one cell following the same order as in D, and each column to a stimulus condition (moving-dots direction or blank, rightmost). Gray lines represent the responses to individual trials, and colored lines represent the median response across trials where the hue corresponds to the preferred direction of the cell. Stimulus presentation periods are shaded red for responses > 1 Z or gray otherwise. Horizontal scale bar: 2 s; vertical scale bar: 5 Z. F. Heatmap of trial-averaged population activity, showing robust responses to motion stimuli but not to the blank condition (rightmost column). Colored bars on the left indicate the tuning type of each cell (yellow: narrow unidirectional, blue: broad unidirectional, red: bidirectional, purple: pandirectional). G. Footprints of cells colored by DSI, showing local clustering of direction selectivity strength. H. Local clustering of direction selectivity, quantified as the mean DSI difference between each cell and its five nearest neighbors, was lower than expected by chance (permutation test, 100 iterations). I. Footprints of cells colored by preferred direction, showing local clustering of directional tuning. J. Preferred direction differences for pairs of neurons sorted by the cell-pair distance. The lower preferred direction differences for cells within ∼200 μm reveals local functional clustering. The oscillatory modulation of preferred direction differences with distance is consistent with a periodic columnar arrangement of direction tuning. The red dashed line at 90° represents the expected random distribution. K. Continuous map of direction selectivity, where each pixel’s color reflects the weighted average DSI of cells within a 200 μm radius. L. Continuous map of preferred direction, where each pixel’s color reflects the weighted average preferred direction of cells within a 200 μm radius. M. Continuous map of local tuning heterogeneity, where each pixel’s brightness reflects the weighted circular variance of preferred directions of cells within a 200 µm radius. Regions with values > 0.9 are highlighted in red, marking putative pinwheel centers and direction fractures.

To test the reliability of each cell’s responses to stimuli, we split the 10 trials for each condition into two sets of 5 trials, then computed the Pearson correlation between the sets. This was then repeated for all possible split permutations (n = 121), resulting in a trial-to-trial reliability index ranging from –1 to 1 for each neuron (STAR Methods). Reliability values close to 1 indicate that the cell’s responses were highly consistent and repeatable across trials, whereas values around 0 indicate no correlation between trials. The average reliability across cells was 0.59 ± 0.25 (median ± SD), indicating highly reliable stimulus-evoked responses to moving dots in MT. To estimate the false positive rate of the statistical tests and thresholds used to detect stimulus-responsive cells, we analyzed ‘responses’ to a blank stimulus during which no stimuli were presented and the gray background kept constant. The blank evoked a mean response larger than 1 Z in only 85 (1.9%) of cells and was the preferred stimulus in none of them, indicating our responsiveness criteria result in a low false-positive rate.

Most cells responded primarily to a few neighboring motion directions, indicating unidirectional tuning. However, we also observed cells responsive to opposite directions along the same axis (‘bidirectional’) and to all directions (‘pandirectional’), as previously reported in macaque MT^77,78^ (**Figure 2F**; **Figure S3**). To classify cells into these three canonical tuning types, we identified for each cell the direction eliciting the maximum response and computed the ortho-index (responses to the ±90° directions divided by the response to the maximally responsive direction) and the anti-index (response to the +180° direction divided by the response to the maximally responsive direction). These indices range from 0 to 1, and correspond respectively to the O/P ratio and 1 – DI used by others^79,80^. Cells with a low anti index were classified as unidirectional, reflecting tuning confined to one direction or a range of neighboring directions. These were further subdivided into narrow unidirectional (low ortho index) or broad unidirectional (high ortho index), depending on whether responses were concentrated at a single direction or widely spread across neighboring directions. Bidirectional cells scored high on the anti-index and low on the ortho index, reflecting two discrete response lobes at opposite directions with suppression at intermediate angles. Pandirectional cells score high on both indices, reflecting approximately uniform responses across all directions. Cells were classified as narrow unidirectional (both indices < 0.5; n = 3,221, 71.7%, tuning width: 52.8 ± 33.3°, median ± SD), broad unidirectional (ortho-index ≥ 0.5, anti-index < 0.5; n = 396, 8.8%, tuning width: 73.5 ± 43.7°), bidirectional (anti-index ≥ 0.5, ortho index < 0.5; n = 611, 13.6%), or pan-directional (both indices ≥ 0.5; n = 264, 5.9%).

As a continuous measure of direction selectivity, we also calculated the direction selectivity index (DSI) of each stimulus-responsive cell as the normalized vector sum of responses across all tested directions (STAR Methods). Cells that responded strongly to one direction and weakly to all others will have a DSI close to 1, whereas cells that responded similarly to all directions will have a DSI close to 0. The DSI across all stimulus-responsive cells was 0.25 ± 0.16 (median ± SD), comparable to previous measurements of inhibitory neurons in marmoset MT^12^ and reflecting strong overall direction selectivity. The relatively large variance in DSI values reflects the diversity of tuning types present in the population, with narrow unidirectional cells exhibiting substantially higher DSI values (median ± SD: 0.385 ± 0.224) than broad unidirectional, bidirectional, and pan-directional cells (0.187 ± 0.110, 0.149 ± 0.101, and 0.102 ± 0.067, respectively).

We then investigated the spatial organization of direction selectivity across MT, asking whether neurons with similar DSIs were spatially clustered (**Figure 2G**). Despite substantial heterogeneity across the recorded FOV, nearby cells tended to have similar DSI values. To quantify this, we computed the mean pairwise DSI difference between each cell and its five nearest neighbors which was significantly lower than the mean of a bootstrapped shuffle distribution (0.16 vs 0.18, respectively, 100 permutations, p < 0.01; **Figure 2H**), indicating that nearby cells are more similar in their direction selectivity than expected by chance. This suggests that the strength of direction selectivity is a locally shared property.

We next characterized the spatial organization of preferred direction across the population. Neighboring cells tended to share similar directional preference, with an arrangement that visual inspection suggests repeats at regular spatial intervals (**Figure 2I**). To quantify this, we calculated the angular difference between the preferred directions for all possible pairs of cells with strong (DSI > 0.3) direction selectivity (**Figure 2J**). Direction differences of 0° indicate tuning to identical directions, values near 180° indicate tuning to opposite directions, and values near 90° indicate orthogonal tuning or, at the population level, a lack of systematic directional alignment. Pairs of cells that were less than 50 µm apart laterally had direction differences of 30.7° ± 49.2° (median ± SD) while pairs that were 200–250 µm apart had direction differences of 97.6° ± 50.3°. Notably, the direction differences did not continue to increase for pairs of cells that were more than 250 µm apart and instead exhibited an oscillatory behavior as a function of the lateral distance between cells. A Fourier analysis of this relationship revealed that dominant spatial frequencies had a period of 400 µm.

Having established features of tuning organization in MT at cellular resolution, we next asked what coarse-grained spatial organization might emerge from such a cellular-level map. While intrinsic imaging generates low-resolution maps where hemodynamic and neuropil signals can obscure single-cell tuning heterogeneity, our mesoscale single-neuron resolution enabled a bottom-up approach, using measurements of individual neurons to construct continuous maps. We first constructed a cortical DSI map by computing the weighted mean DSI of neighboring neurons within a 200 µm radius (STAR Methods). This map revealed how direction selectivity strength varies spatially across MT: with some regions showing locally elevated DSI and others showing lower values, though without a clear global spatial pattern (**Figure 2K**). We next generated a continuous tuning direction map by summing the tuning vectors of neighboring neurons within a 200 µm radius, weighted by each cell’s DSI and distance from the map location (STAR Methods). The resulting map revealed smooth, continuous changes in preferred direction around pinwheel centers across which motion direction jumps rapidly (**Figure 2L**). However, this coarse-grained continuous direction tuning map obscures substantial heterogeneity in single-cell tuning (**Figure 2G, I**) within regions of apparently uniform population preference. Thus, these observations demonstrate that cortical direction maps arise from heterogeneous cellular tuning patterns, providing a direct link between cellular- and large-scale spatial organization.

To investigate whether the observed direction map arises from clustering of like-tuned neurons, we computed a local tuning heterogeneity map (**Figure 2M**). At each location, the local tuning heterogeneity was calculated as the circular variance of the preferred directions of individual neurons within a 200 µm radius, weighted by DSI and distance (STAR Methods). The map values range from 0 to 1, with low values indicating homogeneity and higher values indicating heterogeneity. Local maxima in the map were identified by thresholding at a fixed value of 0.9, allowing detection of pinwheel centers and fractures between neighboring regions exhibiting markedly different directional tuning preferences. These features structurally resemble pinwheels and fractures previously reported in macaque^5^, cat^81^, and ferret^82^ V1, although those were detected by applying a gradient operator to voltage-sensitive dye or intrinsic imaging maps that lack single-cell resolution. In contrast, our local tuning heterogeneity map is constructed directly from the preferred directions and DSIs of individual neurons, revealing additional fine-scale structure, such as continuous spatial gradients and regions of elevated heterogeneity connecting neighboring pinwheel centers.

We repeated these experiments in a second subject, obtaining quantitatively similar results (**Figure S4**). Together, these data confirm the reproducibility of large-scale cellular-resolution imaging in marmosets and the functional organization in marmoset MT, demonstrating how the periodic iso-direction columnar structure arranged into pinwheels and fractures emerge from heterogeneous single cell functional properties.

### Cellular-resolution mesoscopic calcium imaging in marmoset auditory cortex

We next applied our mesoscale cellular resolution 2p calcium imaging platform to marmoset auditory cortex. In primates, the auditory cortex is organized into broad, continuous regions with gradually changing frequency preferences^83,84^, a feature known as tonotopy. In marmosets, tonotopic gradients have been characterized using intrinsic imaging over several square millimeters^85,68^. However, it remains unclear whether the frequency tuning of individual neurons and their neighbors matches that of the larger tonotopic regions in which they are embedded. While electrophysiological experiments show that individual cells can have different tuning than the region in which they are embedded, and nearby cells can be tuned to different frequencies^86^, previous 2p calcium imaging experiments suggest that neural tuning predominantly matches the region and neighboring cells^58^. These experiments used a cytosolically expressed calcium indicator (cyto-GCaMP6f) and did not perform neuropil subtraction, which raises the concern that neuropil-dominated signals may have masked tuning heterogeneity^28,29,87–89^. Revisiting this question, we leveraged the soma-restricted expression of ribo-jGCaMP8m that minimizes neuropil contamination.

We first localized auditory cortex within an optical window with intrinsic imaging during presentation of white noise bursts. Consistent with past studies, regions with strong hemodynamic responses to white noise were predominantly located ventral and posterior to the lateral sulcus (**Figure S3**). We then performed intrinsic imaging during presentation of a series of frequency-ascending and descending tones (STAR Methods). We found distinct areas that predominantly responded to low- or high-frequency tones (**Figure 3A**). Targeting a region that contained both low- and high-frequency responses during intrinsic imaging, we performed cellular resolution single-plane 2p calcium imaging of neurons expressing ribo-jGCaMP8m within a 1.75 × 1.5 mm^2^ FOV (‘FOV1’, **Figure 3A**) at 11.4 Hz. Auditory stimuli consisted of pure tones at seven logarithmically spaced (octave steps) frequencies from 440 to 28,160 Hz, white noise, and marmoset vocalizations (STAR Methods). Among the 2,034 active neurons we detected, 363 (18%) were stimulus responsive. We grouped these cells by the stimulus eliciting their maximal response: 218 (60%) to pure tones, 67 (18.5%) to white noise, 76 (20.9%) to vocalizations, and 2 (0.6%) to stimulus absence. Although 218 neurons showed maximal responses to tones, a larger population (n = 275) responded significantly to at least one tone stimulus (mean Z > 1). We therefore based subsequent analyses on all tone-responsive neurons to study the organization of frequency tuning.

**Figure 3.**
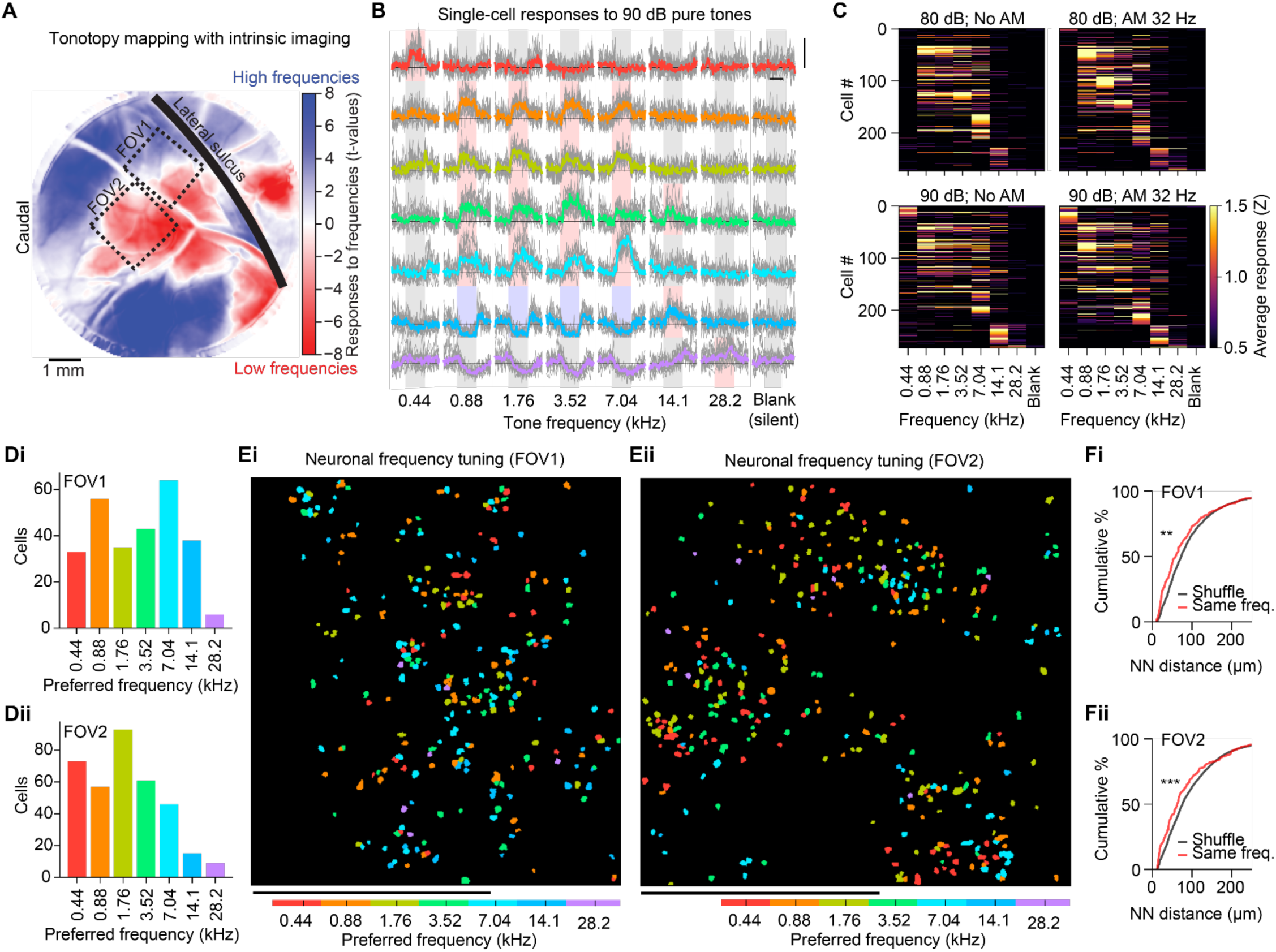
Cellular-resolution mesoscopic imaging and frequency tuning in auditory cortex. A. Functional intrinsic-signal map of an optical window showing regions of auditory cortex that responded predominantly to low-frequency (red) or high-frequency (blue) tones. Two dashed rectangles mark the mesoscopic 2p FOVs shown in panels Ei and Eii. A thick black line indicates the location of the lateral sulcus. B. Stimulus-aligned activity for seven representative cells from FOV1, displaying responses to 90-dB pure tones at seven frequencies (440 Hz to 28.16 kHz). Each row corresponds to one cell and each column to a tone frequency or blank (silent, rightmost). Gray lines represent individual trials and colored lines the median response across trials, where hue corresponds to the preferred frequency of the cell. Stimulus periods are shaded red for responses > 1 Z, blue for responses < −1 Z, or gray otherwise. Horizontal scale bar: 2 s; vertical scale bar: 5 Z. C. Heatmaps of trial-averaged responses of tone-responsive cells from FOV1 across the four stimulus sub-conditions: 80 dB without AM (top left), 80 dB with 32 Hz AM (top right), 90 dB without AM (bottom left), and 90 dB with 32 Hz AM (bottom right). Columns correspond to tone frequencies and blank. D. Number of tone-responsive cells in (i) FOV1, n = 275, or (ii) FOV2, n = 354, that responded maximally to each frequency. E. Spatial distribution of tone-responsive cells in (i) FOV1, n = 275, or (ii) FOV2, n = 354, colored by preferred frequency. Scale bar: 1 mm. F. Cumulative distribution of nearest-neighbor distances for same-frequency cell pairs (red) compared to a bootstrapped shuffle distribution (black) for (i) FOV1 (K–S test: D = 0.12, p < 0.001) or (ii) FOV2.

Responses to auditory stimuli were strong and highly reliable across trials (**Figure 3B**), with a trial reliability of 0.93 ± 0.08 (median ± SD). Most cells exhibited an increase in fluorescence (mean Z ≥ 1), but we also observed suppression (mean Z ≤ −1) in 11 (4%) during stimulus presentation (**Figure 3B**, bottom two rows). While some cells selectively responded to tones of specific frequencies, levels or amplitude modulations, many others responded to combinations of these (**Figure 3C**). To characterize frequency tuning, we identified the frequency eliciting the response maximum, regardless of sound level and amplitude modulation. We found that a similar number of cells preferred each of the presented frequencies except for 28.16 kHz, the highest frequency (**Figure 3Di**). Interestingly, neighboring cells were often most responsive to different frequencies, and the frequency tuning often did not match the regional frequency response pattern seen with intrinsic imaging (**Figure 3Ei**). Nonetheless, neurons preferring the same frequency were located significantly closer to each other than expected by chance (**Figure 3Fi**; median 63 µm vs. 76 µm in shuffle; Kolmogorov–Smirnov test: D = 0.12, p < 0.001). Consistently, the nearest neighbor of each cell shared the same frequency preference in 27.3% of cases, compared to 16.7% in shuffled controls (t-test: p < 0.0001), also supporting modest local clustering.

To test whether this locally heterogeneous organization of frequency tuning extends to other regions of auditory cortex, we recorded from a second FOV of 1.75 × 1.75 mm² at 9.86 Hz in a region identified by intrinsic imaging as biased toward lower frequencies (‘FOV2’, **Figure 3A**). Of 2,873 detected active neurons, 710 (24.7%) responded to an auditory stimulus, with high response reliability (0.89 ± 0.15). We identified 354 cells as tone-responsive, of which 339 (95.8%) showed above-baseline responses, while 11 (3.1%) exhibited suppression and 4 (1.1%) showed mixed responses across tones. Consistent with intrinsic imaging results (**Figure 3A**), FOV2 neurons predominately responded to lower frequencies, with 73 (20.6%) exhibiting a maximum response to 440 Hz compared to only 9 (2.5%) to 28,160 Hz (**Figure 3Dii**). Despite this population-level bias, nearby cells exhibited heterogeneous frequency tuning (**Figure 3Eii**) yet weak clustering (**Figure 3Fii**; Kolmogorov–Smirnov test: D = 0.14, p < 0.0001), mirroring our findings in FOV1. Thus, the combination of single-cell resolution, low neuropil contamination, and large spatial coverage of mesoscopic imaging allowed us to uncover the fine-scale organization of neuronal tuning inside the large-scale maps obtained with intrinsic imaging that mask local heterogeneity.

### Cellular-resolution mesoscopic calcium imaging in the transition area between MT and auditory cortex

Imaging techniques such as intrinsic imaging that include hemodynamic signals as an indirect proxy of neuroactivity, are inherently slow, spatially smoothed, and delayed from the underlying neuronal activity^90–92^. When these techniques describe transitions between functionally different regions as gradients, it remains unclear whether the spatial organization in the transition zone does indeed change gradually or abruptly, only to be blurred in the resulting image.

To characterize neuronal tuning at the boundaries of functionally distinct cortical areas, we leveraged the large FOV of our mesoscope to record from the transition zone between visual and auditory cortices. We identified this region with the intrinsic imaging responses to the previously described moving-dots and white noise stimuli (**Figure 4A**) and then performed single-plane cellular-resolution 2p recordings of neuronal population activity from layer II/III neurons expressing cyto-jGCaMP8s at a depth of 200 µm within an FOV of 2.5 × 2.1 mm^2^ at 2.9 Hz during presentation with stimuli consisting of visual moving-dots in 16 directions, auditory tones at seven octave-spaced frequencies ranging from 440 to 28,160 Hz, marmoset vocalizations, and a blank stimulus (STAR Methods).

**Figure 4.**
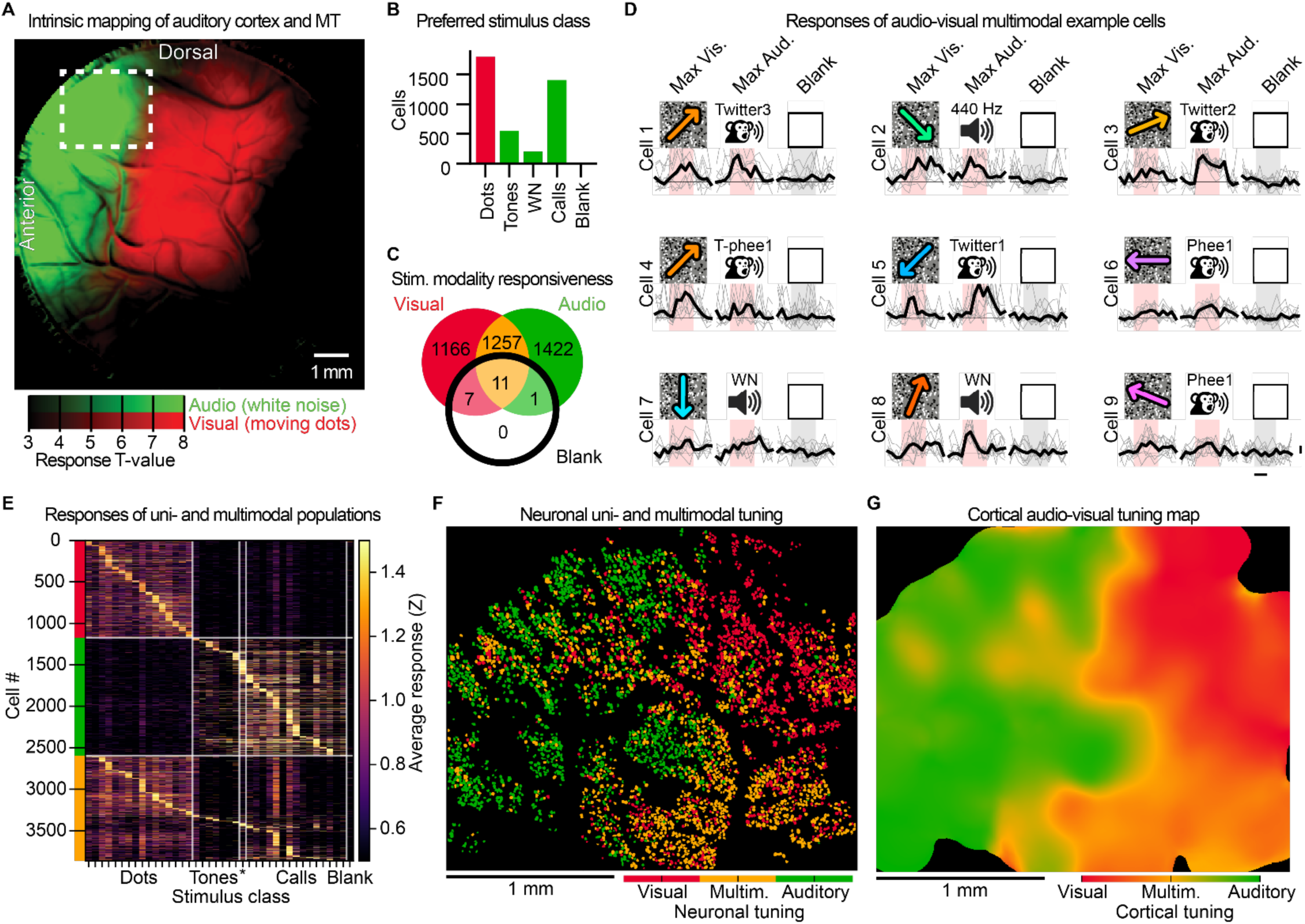
Cellular-resolution mesoscopic imaging of the auditory-visual cortical border and multimodal tuning. A. Functional intrinsic-signal map of an optical window showing adjacent regions with hemodynamic responses to moving dots (red) and white noise bursts (green). The dashed rectangle marks the 2p recording FOV from which the data in panels B–G were obtained. B. Number of cells that responded maximally to each stimulus class: moving dots, pure tones, white noise (WN), marmoset vocalizations (Calls), and blank. C. Venn diagram showing the number of cells that responded significantly (mean Z > 1) to at least one visual stimulus only (red), at least one auditory stimulus only (green), stimuli of both modalities (yellow), and blank (black circle). D. Stimulus-aligned activity for nine multimodal example cells, each shown for three conditions: their maximum visual stimulus (left), maximum auditory stimulus (center), and blank (right). Each column corresponds to one stimulus condition. Gray lines represent individual trials and the black line the median response across trials. Stimulus periods are shaded red for responses > 1 Z or gray otherwise. Auditory stimuli are labeled by call type and caller identity (e.g., Twitter3 refers to a twitter vocalization from animal 3), with five call types each from three different animals. Horizontal scale bar: 1 s; vertical scale bar: 1 Z. E. Heatmap of trial-averaged population responses to moving dots, pure tones, white noise (asterisk), marmoset vocalizations (Calls), and blank. Cells are grouped and sorted by modality preference: visual unimodal (red bar), auditory unimodal (green bar), and multimodal (yellow bar). F. Spatial distribution of cells colored by their modality classification: visual unimodal (red), multimodal (yellow), and auditory unimodal (green). G. Continuous map of audio-visual tuning, where each pixel’s color reflects the weighted average audio-visual tuning index of cells within a 200 µm radius, ranging from visual (red) to multimodal (orange/yellow) to auditory (green).

We detected 3,958 stimulus-responsive cells in our FOV. The largest group, 1,797 (45.4%) cells, preferred visual moving-dots stimuli, followed by 1,407 (35.5%) for vocalizations, 552 (13.9%) for tones, and 199 (5.0%) for white noise, with 3 (0.1%) preferring the blank stimulus (**Figure 4B**). Grouping by modality, 2,158 (54.5%) cells responded maximally to an auditory stimulus and 1,797 (45.4%) to a visual stimulus. This classification is only based on the stimulus eliciting the strongest response and does not imply exclusive selectivity for a single modality. To determine whether cells exhibited responses to both visual and auditory stimuli, we next classified neurons based on whether they significantly responded (mean Z > 1) to at least one stimulus of each modality. We found 1,069 (27.0% of responsive cells) ‘visual unimodal’ cells that responded to at least one visual stimulus but not to any auditory stimulus, 1,516 (38.3%) ‘auditory unimodal’ cells that responded to at least one auditory stimulus but not to any visual stimulus, and 1,373 (34.7%) ‘multimodal’ cells that responded to at least one visual stimuli and at least one auditory stimulus (**Figure 4C–E**). Importantly, the fact that only 19 (0.5%) cells responded with a mean Z > 1 to the blank stimulus, only 3 (0.1%) cells preferred the blank stimulus and no cells responded to the blank stimulus exclusively, are indicative of an overall low false discovery rate.

Across the FOV, auditory unimodal cells were enriched in anterior and dorsal portions, visual unimodal cells were enriched in posterior regions, and multimodal cells were enriched in posterior and ventral portions (**Figure 4F**). Despite these general trends, cells of each response type were also interspersed throughout the recorded region, appearing as isolated individual cells or small local clusters. Notably, this fine-grained heterogeneity in unimodal and multimodal tuning within this transitional region was invisible with intrinsic imaging (**Figure 4A**).

To directly compare the intrinsic imaging map (**Figure 4A**) with the single-cell resolved one (**Figure 4F**), we generated a continuous audio-visual tuning map by assigning each cell an audio-visual tuning index (0 for auditory, 0.5 for multimodal, and 1 for visual; STAR Methods) and summing the tuning indices of all cells within a 200 µm radius, weighted by each cell’s distance from the map location (**Figure 4G**). At the coarse scale, this map replicated the broad posterior-to-anterior transition from visual to auditory cortex seen in intrinsic imaging. However, it also revealed a more complex border structure, with the audio-visual transition following local deviations and irregularities of several hundred microns that were not captured in the intrinsic imaging map.

We performed cellular resolution 2p mesoscopy of the transition zone between MT and the auditory cortex in a second marmoset, obtaining quantitatively similar results (**Figure S5**). Taken together, these results demonstrate how large-scale cellular resolution mesoscopy can resolve cellular tuning and spatial intermingling at the boundaries of large cortical territories. Notably, overlapping signals in intrinsic imaging could reflect either true multimodal responses, a mix of unimodal responses, or merely the spatial blurring across neighboring unimodal regions. In this context we demonstrate how our large-scale single-cell neuronal recording capability reveals the presence of both unimodal and multimodal cells and that functional boundaries between cortical areas are more complex and variable than what intrinsic imaging alone or what simple gradient model would suggest. Thus, cellular resolution at a mesoscopic scale is necessary to resolve this organization.

### Volumetric mesoscopic recording of neuronal population activity with cellular resolution in marmoset area MT

Cortical neuronal population do not only exhibit organization along the lateral directions that form various cortical regions but also along the axial direction as evidenced by the canonical columnar organization of the cortex with its layer-specific functional and anatomical connections. Mesoscopic single-plane recordings cannot capture neuronal tuning along the axial dimension and constrain the total number of simultaneously recorded neurons. Extending point-scanning imaging methods to three-dimensional (3-D) volumetric recordings has been conventionally limited by the required temporal tradeoff that results in prohibitively slow volume rates.

Light Beads Microscopy^11^ (LBM) has been shown to overcome this issue by capturing activity across 30 axially separated excitations along cortical depths within ∼200 ns and thereby enable mesoscopic volumetric cellular-resolution calcium imaging of up to a million neurons distributed across both hemispheres and different layers of the mouse cortex at multi-Hertz rate^11,14^. Realizing such large neuronal population recordings in the scattering mammalian brain requires a maximally efficient and scalable acquisition scheme that optimally navigates the tradeoffs between acquisition speed, scale of the recording, resolution and photon-efficient generation of fluorescence within the safe limits of power exposure of the sample. LBM uniquely addresses these requirements by ensuring that: (i) each sample voxel is excited by a single laser pulse, (ii) spatial sampling is matched to the size of the structures of interest, and (iii) temporally adjacent voxel acquisitions are separated by no more than the fluorescence lifetime^11^. LBM achieves all of the above criteria simultaneously via a cavity-based 30-fold spatiotemporal multiplexing utilizing the Many-fold Axial Multiplexing Module (MAxiMuM) ^11^. MAxiMuM converts each high-energy femtosecond laser pulse into a set of axially separated foci (“light beads”) that are temporally spaced by the fluorescence lifetime and distributed across the full axial imaging range (∼500 µm) which is acquired within ∼ 200 ns. This results in a fluorescence-lifetime limited pixel acquisition rate of ∼150 MHz which has amongst different configurations enabled cellular resolution volumetric calcium imaging within volumes of ∼6.0 × 5.4 × 0.5 mm^3^ at ∼2.2 Hz in mice. Additionally, the modular design of MAxiMuM allows flexible multiplexing configurations, including adjustments to the number of light beads, per-bead energy density, axial bead separation, total axial coverage, and inter-bead power falloff, making it adaptable to the specific optical constraints of different biological requirements.

To obtain an in-depth characterization of MT’s functional organization and to increase neuronal yield we performed cellular-resolution mesoscopic scale simultaneous recording across cortical depths using LBM in the marmoset. Since marmoset cortical layer I is largely devoid of cell bodies and thicker than in mice, we configured LBM for mesoscale volumetric recordings only of the neuron-dense layer II/III, using 15 light beads each axially separated by 20 µm, collectively spanning a 300 µm axial imaging range. The MAxiMuM cavity allowed for an exponential increase of the pulse energy of the light beads as a function of depth that was matched to the scattering length of the tissue and allowed to compensate for scattering-induced attenuation and to maintain constant SNR across depth (**Figure 5A**). Total average power was kept below 300 mW, and lateral power density kept below 60 mW/mm², well below the established safety thresholds for 2p imaging^70,21,22,11^. In addition, using this configuration with only 15 light beads allowed to effectively eliminate the temporal cross talk between temporally-adjacent light beads by increasing their temporal separation to ∼15 ns. Under these conditions, we performed an LBM recording in the central region of MT within a volume of 2 × 2 × 0.3 mm^3^ spanning 100–400 µm depths at 2.6 Hz in an FOV expressing soma-jGCaMP8s while presenting moving-dots stimuli in 16 directions. To prevent redundant detection of the same neuron across adjacent planes, segmented cell ROIs were axially merged when they were sufficiently close (<10 µm lateral displacement, ≤4 planes) and exhibited correlated (Pearson *r* > 0.15) fluorescence activity (STAR Methods). All other quantifications were performed identically to those used for single-plane recordings.

**Figure 5.**
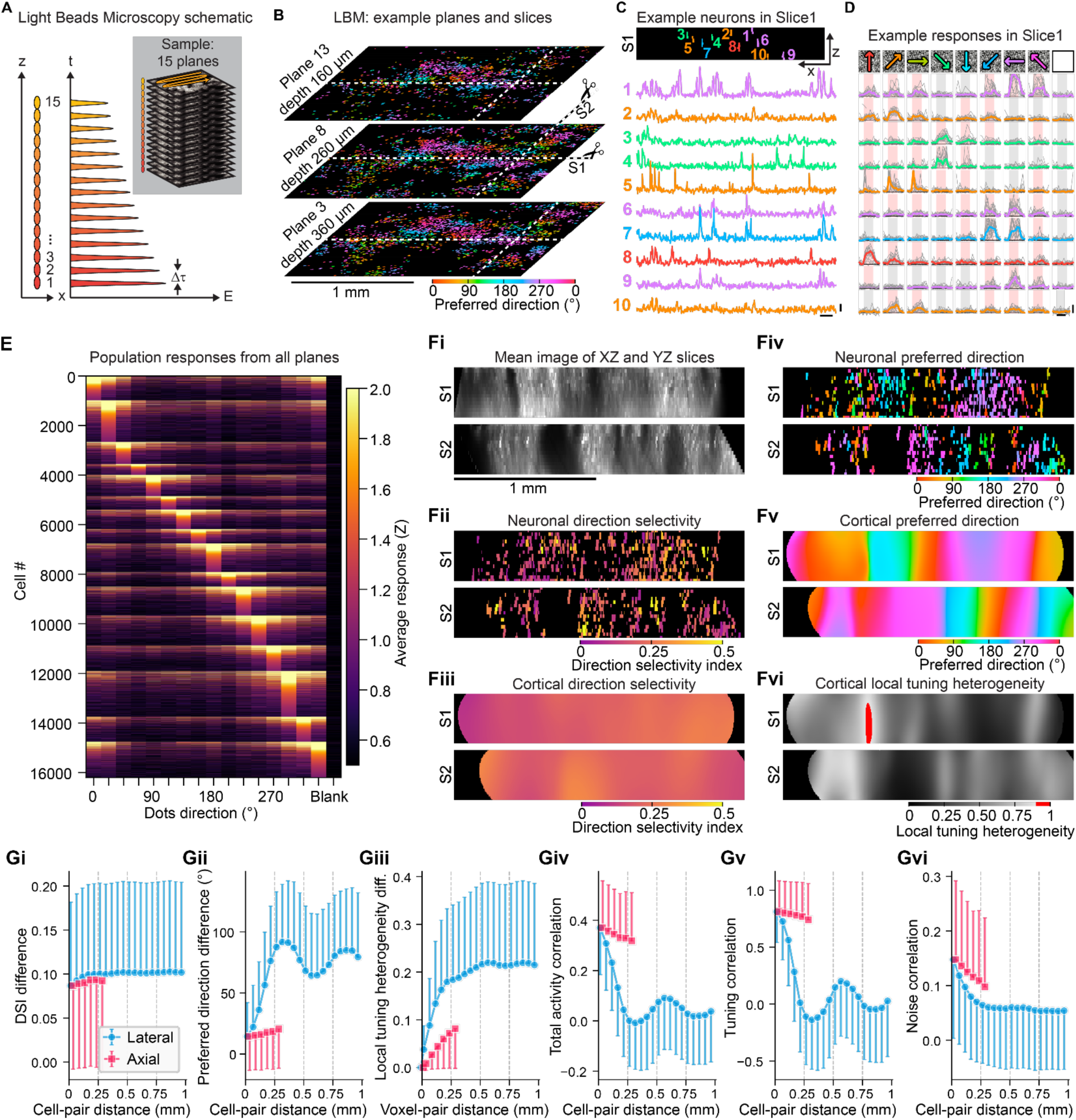
Volumetric functional characterization of direction tuning in MT using Light Beads Microscopy (LBM). A. We used LBM to split each laser pulse into 15 copies that focus along an axial column of ∼300 μm enabling near-simultaneous volumetric imaging. To maintain SNR across the volume, deeper beads carry higher pulse energies to account for increased scattering. B. Three of the 15 planes imaged with LBM, each of 2 × 2 mm², with cells colored by preferred direction. Dashed lines indicate the Y position for the XZ slice (S1) and the X position for the YZ slice (S2) shown in 5C and 5F. C. The top panel shows an orthogonal XZ view (S1) from the recording volume where 10 footprints of selected example cells were colored by their preferred direction (color conventions as in B). The bottom panel shows the activity traces of the example neurons. Horizontal scale bar: 10 s; vertical scale bar: 5 Z. D. Stimulus-aligned activity for the 10 cells shown in panel 5C displays stimulus-locked responses to different subsets of moving-dots directions. Each row corresponds to one cell following the same order as in 5C, and each column to a stimulus condition (moving-dots direction or blank, rightmost). Gray lines represent the responses to individual trials, and colored lines represent the median response across trials where the hue corresponds to the preferred direction of the cell. Stimulus presentation periods are shaded red for responses > 1 Z or gray otherwise. Horizontal scale bar: 2 s; vertical scale bar: 5 Z. E. Heatmap of trial-averaged population activity, showing strong responses to motion stimuli but not to the blank condition (rightmost column). F. Orthogonal views of the slices S1 and S2 showing: (i) mean image; (ii) neuronal DSI; (iii) cortical DSI map; (iv) neuronal preferred direction; (v) continuous map of preferred direction; and (vi) local tuning heterogeneity map. G. Pairwise-distance analysis across the volume (median ± SD) for: (i) neuronal DSI difference, (ii) neuronal preferred-direction difference, (iii) cortical local tuning heterogeneity difference across voxels, (iv) neuronal total activity correlation, (v) tuning correlation, and (vi) noise correlation. Data corresponding to lateral (XY, blue) and axial (Z, red) separations are shown separately. For each datapoint, the SD is shown only in one direction to aid visualization.

Neurons exhibiting strong fluorescence dynamics and responses to stimuli were present across all imaging planes (**Figure 5B–D, Video S4**). Within the recorded volume, we detected 39,964 active neurons of which 16,205 (40.5%) responded to moving-dot stimuli (**Figure 5E**). The analysis of individual planes within our recording volume revealed comparable fluorescence traces, robust stimulus-driven responses, and similar numbers of responsive neurons across depths (**Figure S6A–D**). Cells had a response reliability of 0.68 ± 0.22, indicating reproducible responses across trials and consistent with the observed values in our single-plane recordings for animals expressing ribo-jGCaMP8s, suggesting a similar SNR across imaging approaches.

We next investigated the 3-D spatial organization of the strength of direction tuning. Cells had a mean DSI of 0.19 ± 0.11, and cells with similar DSIs appeared to show weak spatial clustering along both lateral and axial dimensions (**Figure 5Fi–Fii**). We computed a 3-D continuous DSI map (**Figure 5Fiii**), with no clear spatial pattern across the volume. To quantify potential spatial clustering of direction selectivity, we computed the difference in DSI between all neuron pairs as a function of the lateral (XY) and axial (Z, i.e. depth) distance between cell bodies (**Figure 5Gi**). These results reveal weak local clustering of cells with similar DSI within ∼200 µm, but no systematic lateral or axial dependence and no clear spatial pattern at larger distances, indicating an absence of clear columnar structure for direction selectivity magnitude.

In contrast, cells with similar preferred directions exhibited stronger spatial clustering throughout the recorded volume (**Figure 5Fiv, Video S5**). The continuous map of preferred direction revealed that iso-direction regions spanned vertically throughout the recorded volume (**Figure 5Fv**). When analyzing the difference in preferred direction for pairs of neurons, cells that were separated by less than 50 µm either laterally or axially had a median difference in preferred direction of 14 ± 28° (**Figure 5Gii**). For neuron pairs separated by 250–300 µm axially but less than 50 µm laterally, this value was slightly larger at 21 ± 32°. In contrast, for neuron pairs separated by 250–300 µm laterally but less than 50 µm axially, the median difference in preferred direction was markedly higher at 88 ± 45°, reaching a global maximum. These results indicate that direction tuning is organized anisotropically, with lateral separation impacting tuning dissimilarity more strongly than axial separation, consistent with a columnar organization of direction tuning in MT. At lateral separations beyond 250–300 µm, preferred direction differences exhibited an oscillatory dependence on distance with a period of ∼500 µm, consistent with our single-plane observations with ribo-jGCaMP8s (**Figure 2, Figure S4**).

We next generated 3-D local tuning heterogeneity maps (**Figure 5Fvi**). We found that regions maintained similar values throughout the recorded depth. Since local tuning heterogeneity is not a neuronal property but an emergent property of a cortical region, we quantified local tuning heterogeneity differences as a function of lateral and axial voxel-pair separation (**Figure 5Giii**). For voxel pairs separated by less than 50 µm laterally or axially, local tuning heterogeneity was 0.04 ± 0.05 laterally and 0.01 ± 0.01 axially. For voxel pairs separated by 250–300 µm laterally, this value was 0.18 ± 0.16, and for pairs separated by 250–300 µm axially, this value was 0.08 ± 0.08. The faster rate of change along the lateral dimension indicates that regions of heterogeneous direction tuning are organized anisotropically, with greater conservation along the axial dimension, consistent with pinwheel centers and direction fractures spanning vertically across cortical depth. Interestingly, cells in areas of high local tuning heterogeneity tended to have lower DSIs, suggesting that regions of diverse directional preferences are populated by less selective neurons. While this relationship was highly significant (linear regression, p < 10^-10^), the effect size was modest (R² = 0.10), indicating that local tuning heterogeneity explains only a small fraction of the variance in individual cell DSI. Thus, while the trend is robust and consistent across the population, local tuning heterogeneity alone is not a strong predictor of single-cell direction selectivity.

While direction tuning can be estimated from sequential recordings of large neuronal populations, only simultaneous volumetric imaging enables functional correlation analysis across both lateral and axial dimensions at single-cell and single-frame resolution. Such analysis is essential because pairwise activity correlations between any two neurons can arise from at least two distinct sources: similarity in stimulus tuning, which produces correlated trial-averaged responses (tuning correlation), and trial-to-trial covariability reflecting direct or indirect local connectivity or common upstream inputs (noise correlation)^93,94^. Disentangling these contributions is critical for determining whether the spatial pattern of activity correlations reflects direction tuning organization or local circuit connectivity^95^. We leveraged the simultaneous nature of LBM volumetric recordings to compute correlations in the fluorescence signals for all neuron pairs as Pearson correlation coefficients. Activity correlations were highest for nearby neurons, with a median of 0.37 ± 0.19 for pairs separated by less than 50 µm laterally or axially (**Figure 5Giv**). Along the axial dimension, correlations declined only modestly, remaining at 0.33 ± 0.19 for pairs separated by 250–300 µm. In contrast, activity correlations decreased rapidly with increasing lateral separation, dropping to 0.00 ± 0.18 for pairs separated by 250–300 µm laterally, and then exhibited an oscillatory pattern with a spatial periodicity of ∼500 µm. Notably, the median activity correlation remained positive for all lateral and axial distances sampled.

The period pattern of correlations (**Figure 5Giv**) could be due to correlations of tuning preference (**Figure 5Gii**), which exhibits a very similar pattern, or due to noise correlations or both. To separately quantify tuning and noise correlations, we decomposed each neuron’s single-trial response into the sum of its trial-averaged response to that stimulus and a trial-specific residual^94,95^. Tuning correlations were then computed as the correlation between the trial-averaged responses of each neuron pair, whereas noise correlations were computed as the correlation between their trial-specific residuals. Tuning correlations for neurons less than 50 µm apart laterally and axially were 0.81 ± 0.27, were high for cells axially separated by 250–300 µm (0.74 ± 0.32) but sharply lower for cells separated by 250–300 µm laterally (−0.11 ± 0.42). Beyond this distance, tuning correlations exhibited an oscillatory dependence on lateral separation with a periodicity of ∼500 µm. Noise correlations were 0.15 ± 0.14 for nearby cells (less than 50 µm laterally or axially) and of comparable magnitude along both lateral (0.06 ± 0.11) and axial (0.10 ± 0.13) dimensions at 250–300 µm separation, indicating an approximately isotropic decay. Beyond 300 µm of lateral separation, noise correlations decayed monotonically, though still positive for lateral distances of up to 2 mm. These results indicate that while neurons within a given column exhibit substantial shared trial-to-trial variability, columns with similar direction tuning do not show elevated noise correlations compared with columns tuned to other directions. Thus, shared trial-specific variability depends primarily on proximity rather than on the spatial organization of direction tuning.

To capture as much of the MT population as possible, we next extended the spatial range of volumetric imaging to cover a FOV of 4 × 4 × 0.3 mm³ by relaxing pixel spacing to 3.5 µm which was recorded at 1.9 Hz (**Figure S7, Video S6–S8**). Considering a median neuron diameter of 13 µm in marmoset cortex^96^, our relaxed pixel spacing of 3.5 µm allows each neuron to be sampled by ∼14 pixels, sufficient to resolve individual neurons and consistent with previous results in which it has been shown that even a higher pixel spacing of 5 µm is still compatible with single-cell detection^11^. Within this larger FOV, which combined soma- and cyto-jGCaMP8s expressing regions, we could capture 96,828 active neurons, of which 46,126 (47.6%) responded to moving dots and moving gratings in 8 directions, while yielding similar DSI, functional organization of direction tuning, and signal correlations as observed in our smaller FOV recordings. Finally, the above experiments were repeated in a second marmoset, obtaining quantitatively similar results (**Figure S8**). Together, these findings confirm the reproducibility of LBM imaging in marmoset cortex and the ability to reveal 3-D spatial organization of functional properties in very large populations of cells.

Overall, these results demonstrate how mesoscale cellular resolution volumetric recording capabilities of LBM enable a 3-D characterization of the spatial organization of neuronal tuning and functional correlations in the marmoset cortex. By simultaneously recording neuronal population activity across large laterally extended FOVs and axially along depth in marmoset MT, LBM revealed a columnar organization of direction tuning across cortical depth and a periodic spatial organization laterally. Moreover, LBM’s simultaneous volumetric imaging capability enabled a functional dissection of activity correlations, revealing that tuning correlations reflect the spatial organization of direction preference, whereas noise correlations are governed primarily by physical proximity rather than by shared tuning. Together, these findings establish LBM imaging as a powerful tool for linking 3-D functional maps to the underlying structure of neuronal tuning and connectivity in NHPs.

### Properties of direction tuning across the MT neuronal population

Previous studies of direction tuning in macaque MT qualitatively classified cells based on their direction selectivity, finding most neurons (∼85%) to have unidirectional tuning, and smaller fractions of bidirectional (7%) and pandirectional (8%) cells^77,78^. While visual inspection of the tuning curves and classification might be feasible when only a few hundred cells are recorded, this approach is not scalable to larger neuronal recordings where more rigorous quantitative methods are required. Moreover, the bidirectional and pandirectional populations may have been underrepresented in past recordings due to their lower prevalence and the limitations of single-cell sampling. Additionally, it remains unclear whether cells with distinct direction tuning profiles reflect discrete functional subpopulations or continuously vary along a tuning spectrum.

To characterize the properties of motion tuning across the population in a data-driven manner, we performed principal component analysis (PCA) on the neuronal responses to moving dots stimuli from our largest MT recording (4 × 4 × 0.3 mm³ FOV at 1.9 Hz; **Figure S7**). We restricted this analysis to neurons with a mean activity of Z > 1.5 for each cell’s maximally responsive direction to ensure reliable estimation of responses across all other directions, including 16,614 (41.2% of dots-responsive) cells. Each neuron’s response was represented as a 16-dimensional tuning vector of trial-averaged responses to each motion direction, yielding a 16,614 × 16 (neuron × direction) matrix. PCA decomposed this matrix into 16 principal components (PCs), each a 16-dimensional vector of loadings across motion directions, ordered by the variance they explained in population responses (**Figure 6A**). The first five PCs accounted for 66.2% of total variance (PC1: 19.1%, PC2: 17.1%, PC3: 15.2%, PC4: 7.7%, PC5: 7.1%; **Figure 6B**). Each of these first five PCs captured a distinct aspect of directional tuning (**Figure 6C**). PC1 had concentrated positive loadings in the downward direction and negative loadings in the upward direction, while PC2 had concentrated positive loadings in the left direction and negative loadings in the right direction. Since individual cells can have either positive or negative weights for each PC, PC1 and PC2 together forming a basis for unidirectional tuning. PC3 had approximately uniform loadings across all directions, matching the expected tuning of pandirectional cells. Analogously to how PC1 and PC2 together span all possible unidirectional tuning preferences and formed a basis for unidirectional tuning, PC4 and PC5 each showed two prominent opposing loading concentrations, with the main axes of the two PCs offset by 45° from each other. Because individual cells can have either positive or negative weights for each PC, any bidirectional tuning curve, regardless of its preferred axis, can be expressed as a linear combination of PC4 and PC5, and thus represent a basis for bidirectional tuning.

**Figure 6.**
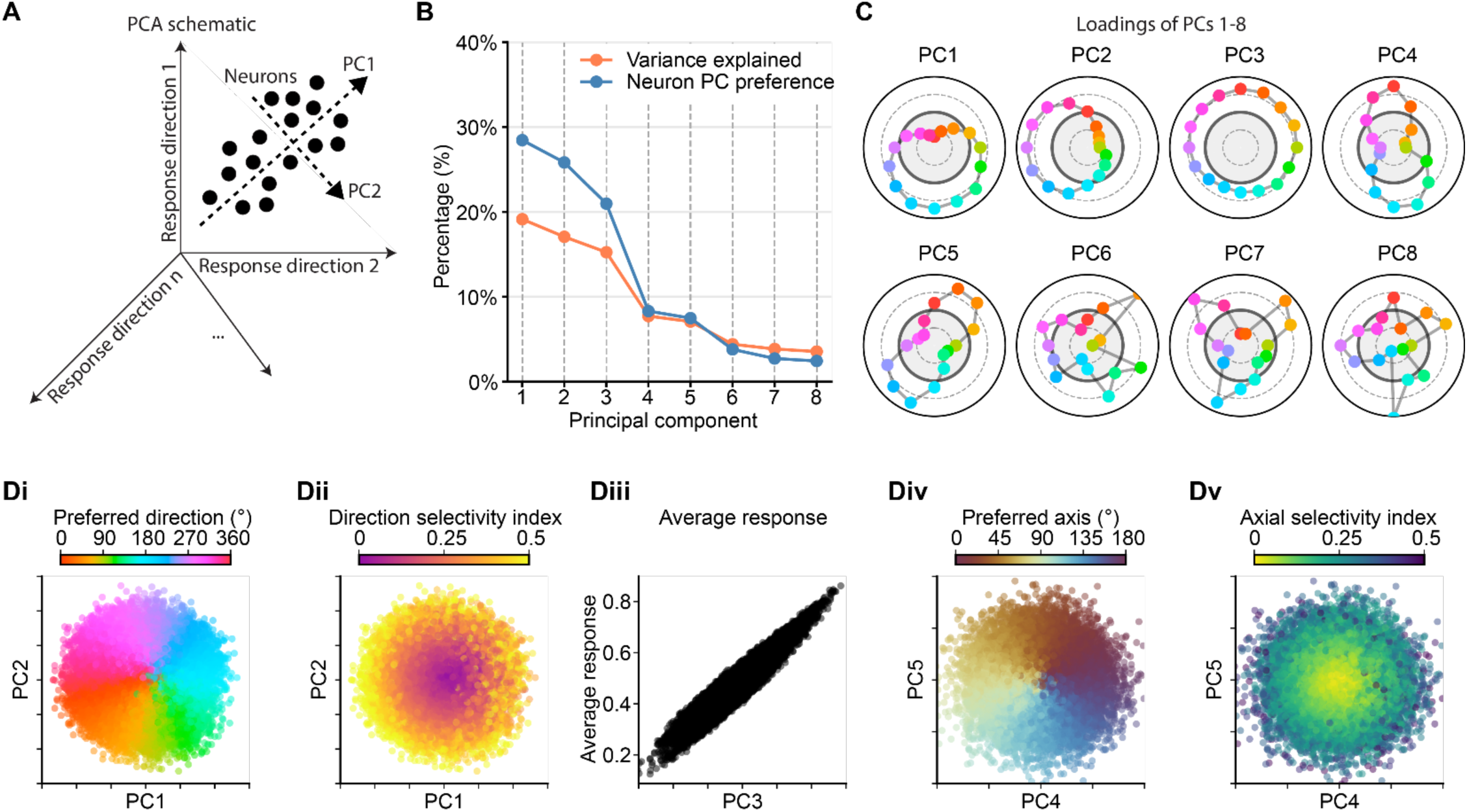
Population analysis of motion encoding in MT. A. Schematic of PCA, showing how PCs capture variance across neurons in the space generated by responses to the 16 motion directions. B. Explained variance (orange) and neuron PC preference (blue) for each of the first eight PCs. Neuron PC preference was defined as the component with the largest absolute loading for each cell. C. PC loadings across the 16 motion directions (points colored by direction) for the first eight PCs. PC1 and PC2 capture unidirectional tuning, PC3 uniformly captures responses across all directions, and PC4 and PC5 capture bidirectional tuning. D. Single-cell projections onto selected PC pairs, colored by independently derived tuning metrics. (i) PC1 versus PC2, colored by preferred direction of motion. (ii) PC1 versus PC2, colored by DSI. (iii) PC3 versus average response amplitude. (iv) PC4 versus PC5, colored by preferred axis of motion. (v) PC4 versus PC5, colored by ASI.

We next asked whether these PCs aligned with individual neuronal tuning profiles by identifying the PC with the largest absolute contribution ("preferred PC") for each neuron. If the loadings of individual PCs did not capture the tuning profiles of distinct functional cell types, the fraction of cells preferring each PC would simply reflect its share of explained variance. Instead, we found that the first five PCs aligned with the tuning of disproportionately more cells than their explained variance would predict, while higher PCs aligned with that of very few neurons (**Figure 6B**). We therefore collapsed PC1 and PC2 into a single unidirectional subspace and PC4 and PC5 into a single bidirectional subspace based on their joint contributions (STAR Methods). This concentrated cell assignments into three representational bases: unidirectional (PC1–PC2: 55.4%), pandirectional (PC3: 16.9%), and bidirectional (PC4–PC5: 15.7%), with the remaining 6.0% distributed across all other PCs, indicating that no other functional tuning groups were substantially represented in the population.

We then examined how direction tuning was encoded in low-dimensional PC spaces by linking PC scores and stimulus responses for each neuron. In the PC1–PC2 plane, cells were arranged in a polar coordinate system in which preferred direction determined angular position and DSI determined radial distance, with strongly unidirectional cells displaced toward the periphery (**Figure 6Di–ii**). PC3 was strongly positively correlated with mean response amplitude (R² = 0.906, p < 10⁻^10^; **Figure 6Diii**) and negatively correlated with DSI (R² = 0.322, p < 10⁻^10^; **Figure S9Bii**), consistent with its distributed loadings across all directions. To characterize bidirectional tuning, we computed a preferred axis and an axial selectivity index (ASI) by effectively folding the 360° direction space onto a 180° axis space and computing the vector sum of responses (STAR Methods). Analogously to the representation of unidirectional tuning in PC1–PC2, cells were arranged in the PC4–PC5 plane in a polar coordinate system in which preferred axis determined angular position and ASI determined radial distance, with strongly bidirectional cells displaced toward the periphery (**Figure 6Div–v**).

Together, these analyses demonstrate that the direction-tuning properties of MT neurons are well captured by a low-dimensional population subspace, with the first five PCs forming interpretable bases for the three observed tuning classes, collectively accounting for most of the response variance across the MT population. Further, and despite previous studies classifying as separate and distinct functional types, the distribution of neurons within low-dimensional PC subspaces appeared continuous, with cells spanning the entire PC space rather than forming discrete clusters and suggesting a functional continuum.

### Functional diversity and spatial organization of direction tuning properties in MT

These results together with the very large size of the neural population we measured, open the possibility to address the key question of whether the continuous distribution of neurons within low-dimensional PC subspaces reflects genuine variation in single-neuron tuning profiles, or whether it is an artifact of PCA, which is optimized to decompose population-level variance and does not necessarily correspond to the tuning profiles of individual cells. To address this, we utilized the previously introduced ortho- and anti-indices, which provide a continuous, physiologically interpretable characterization of each cell’s direction tuning profile.

For each neuron, the ortho-index and anti-index were computed using a cross-validated three-way trial split: the first third of trials were used to identify the maximally responsive direction, the second third to compute the indices, and the final third to estimate the full tuning curve in reference to the direction identified by the first split. This was repeated 100 times with randomly drawn trial splits, and the resulting indices and tuning curves were averaged across repetitions to yield cross-validated estimates for each cell.

The joint distribution of ortho and anti-indices confirmed the presence of all four tuning types (**Figure 7A**). Applying an arbitrary 0.5 threshold to the indices as done for our single-plane recordings, we identified 7,488 (45.1%) cells with narrow unidirectional tuning, 3,710 (22.3%) with broad unidirectional tuning, 1,490 (9.0%) with bidirectional tuning, and 3,926 (23.6%) with pandirectional tuning. Critically, however, the distribution across this 2-D space revealed no discrete clusters or abrupt transitions between groups. Instead, cells densely tiled the entire anti index and ortho index continuum.

**Figure 7.**
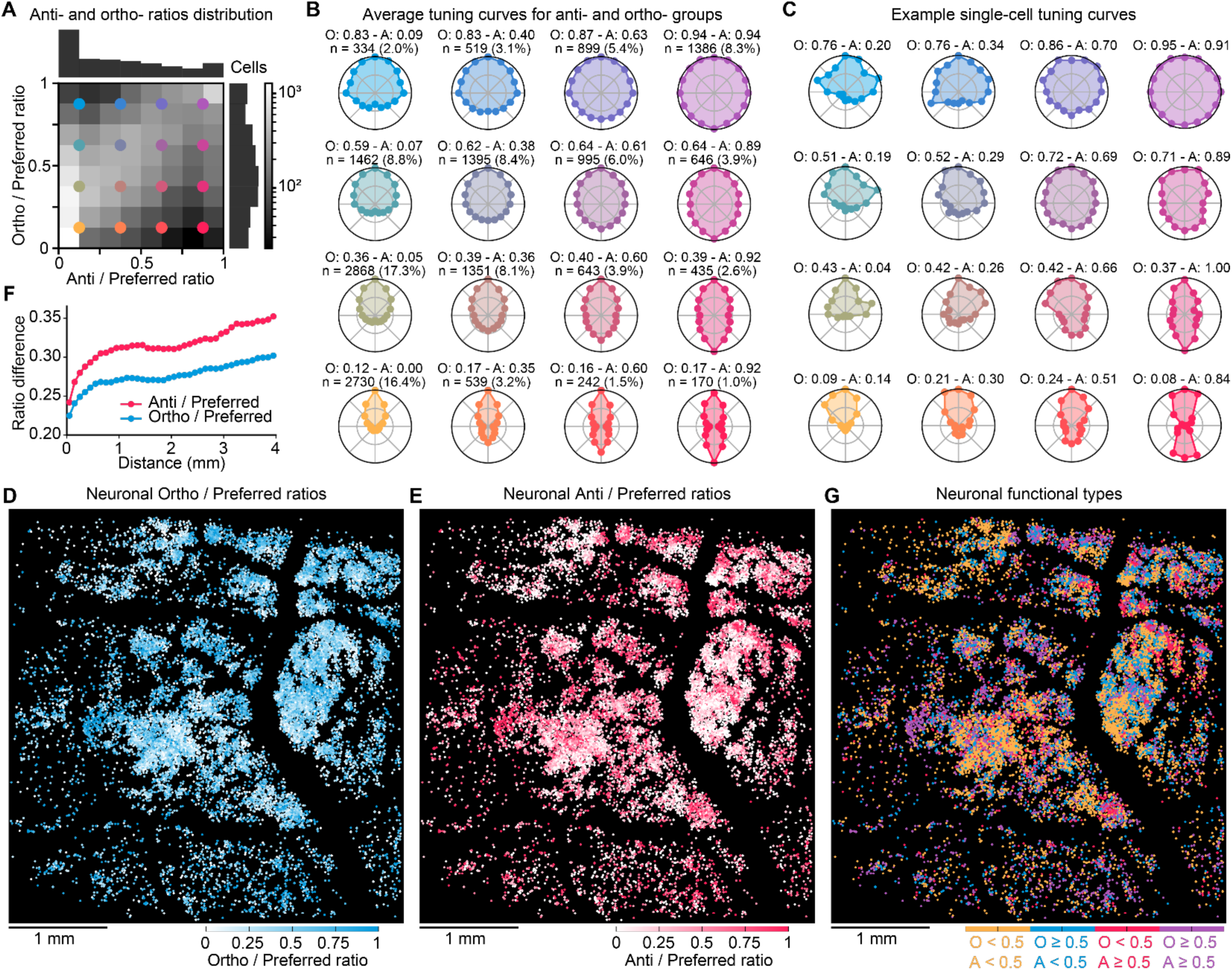
Functional diversity and spatial organization of direction tuning properties in marmoset MT. A. Joint distribution of anti index (A) and ortho index (O) across the MT population, displayed as a 2D histogram with marginal distributions. Each index is computed relative to the preferred response. Colored markers indicate the centers of the 4×4 A/O bins used for grouped tuning summaries in B and C. B. Population-average direction tuning curves arranged in a 4×4 grid of A/O bins. Each polar plot shows the mean normalized tuning curve of all cells assigned to that bin. Subplot titles report the bin cell count (n), the median ortho index (O), and the median anti index (A) across cells in the bin. C. Representative single-cell tuning examples arranged in the same 4×4 A/O grid as B. Each polar plot shows one example neuron from the corresponding bin, illustrating how individual response properties relate to the bin-average in B. Subplot titles report the cell’s own ortho index (O) and anti index (A), each computed as the median across 100 trial splits. D. Spatial map of ortho index. Direction-selective cells across all 15 LBM planes were projected onto a common XY plane and colored by their ortho index using a white-to-blue scale. E. Spatial map of anti index. The same projection as D, with cells colored by their anti index using a white-to-red scale. F. Distance dependence of tuning dissimilarity. Median absolute pairwise difference in ortho index (blue) and anti index (red) is plotted as a function of cell-pair distance (100 µm bins), revealing that nearby cells are more similar in their tuning properties than distant ones. G. Spatial map of functional tuning classes. Cells are classified by applying a 0.5 threshold jointly to both the anti index and ortho index, yielding four types: narrow unidirectional (A < 0.5, O < 0.5; yellow), broad unidirectional (A < 0.5, O ≥ 0.5; blue), bidirectional (A ≥ 0.5, O < 0.5; orange), and pandirectional (A ≥ 0.5, O ≥ 0.5; purple).

To test whether cells occupying intermediate regions of this space represent genuine intermediate tuning profiles rather than misclassifications, we subdivided the index space into a 4 × 4 grid and computed the mean tuning curve for each bin. Tuning curves in intermediate bins smoothly varied between those of the extreme groups across all combinations of indices (**Figure 7B**), and example cells within each bin closely matched their subgroup (**Figure 7C**). This establishes that intermediate cells reflect a true continuum of response profiles rather than a mixture of extreme tuning types. Together, these single-cell results confirm that the continuity observed in the PC subspace is not a mathematical artifact of the population decomposition, instead reflecting genuine variation in the tuning properties of individual neurons. Marmoset MT thus contains a continuous spectrum of direction tuning, with cells smoothly spanning the space between extremes.

The existence of patterns in the distribution of ortho and anti-tuning across MT would indicate functional organization of the unidirectional, bidirectional, and pandirectional tuning classes. Therefore, we next asked whether ortho and anti-tuning was spatially organized across MT. Both the ortho index (**Figure 7D**) and anti-index (**Figure 7E**) exhibited significant local clustering. Nearby neurons shared more similar values than distant pairs and pairwise differences increased monotonically with distance, with ortho indices changing more rapidly with distance than anti indices (**Figure 7F**). At larger spatial scales, neither index exhibited a clear periodic structure, in contrast to the ∼500 µm periodicity observed for preferred direction (c.f. **Figure 5Gii**). This suggests that cells with similar tuning profiles aggregate locally but local clusters are not arranged in a repeated columnar pattern. Assigning cells to the four tuning types using the 0.5 threshold confirmed local clustering of cells with similar tuning profiles across MT (**Figure 7G**).

Finally, we investigated whether cells with distinct tuning properties exhibited differential selectivity for a second stimulus class: moving sinusoidal gratings (**Figure S10**). Of the 16,614 cells that exceeded the Z > 1.5 threshold for moving-dots stimuli, 6,661 (40.1%) also responded with Z > 1.5 to at least one grating direction. However, the probability of responding to gratings varied systematically across the anti-index and ortho-index space. Cells with strongly narrow unidirectional tuning (both indices < 0.25) were least likely to respond (31.5%), while cells with strongly bidirectional tuning (anti index > 0.75, ortho index < 0.25) were most likely (60.6%).

For the subset of cells that responded to both dots and gratings, the relationship between preferred dot direction and preferred grating direction also differed across classes. Narrow unidirectional cells tended to respond to gratings moving in the same direction as their preferred dot motion (**Figure S10**, bottom left), consistent with encoding the direction of motion regardless of stimulus class. In contrast, bidirectional cells tended to respond to gratings moving orthogonally to their preferred dot direction, consistent with their tuning to the axis of motion matching the spatial orientation of the grating (**Figure S10**, bottom right). Taken together, these results indicate that cells with distinct direction tuning profiles differ not only in how they respond to dot motion, but also in their selectivity for a qualitatively different stimulus and the features they extract from it, suggesting different motion encoding strategies.

## DISCUSSION

Neuroimaging technologies are constrained by inherent tradeoffs between scale of recording, speed, resolution and neuronal SNR. As a result, most existing techniques due to their suboptimal design lack the ability to record simultaneous large-scale, volumetric, multi-regional neuronal population activity at cellular resolution at high speed in the mammalian brain. While significant advances have been recently achieved through the combined development of cellular-resolution mesoscopy platforms ^10,71,23^ and highly optimized 2p acquisition approaches^11^, major challenges have prevented the extension and application of these technologies in NHPs.

Here, we present the development and application of an optimized 2p calcium imaging platform for NHPs that integrates mesoscale, cellular-resolution volumetric recording of large neuronal population activity with improved calcium indicators that have been optimized for marmosets. By combining calcium indicator expression systems tailored for the primate brain with a mesoscopy platform and LBM implementation adapted for NHP brain imaging, we achieved volumetric neuronal population recordings in visual and auditory areas spanning up to ∼4 × 4 × 0.3 mm³ at single-cell resolution in awake marmosets (**Figure S7**). These recordings simultaneously captured the activity of over 96,000 active cells—including over 46,000 stimulus-responsive cells—at 1.9 Hz, which to our knowledge represents by far the largest simultaneous recording of cellular-resolution neuronal population activity in a primate to date.

By spanning micrometric spatial resolution across millimeter-scale FOVs, this technique provides, for the first time, a unified experimental framework for directly linking neuronal tuning properties with large-scale functional organization in NHPs. This large-scale, cellular-resolution approach contrasts with the two-stage experimental strategy typically applied in NHP research: low-resolution, large-scale methods such as intrinsic imaging, fMRI, and one-photon imaging to identify millimeter-scale functional areas, followed by high-resolution, small-scale methods such as 2p imaging or electrophysiology to record individual neuronal responses. While this classical strategy has substantially advanced our understanding of primate cortical organization, it implicitly assumes that neurons sampled within restricted regions are representative of entire functional areas. Consequently, tuning properties measured from tens to hundreds of neurons are extrapolated across millimeter-scale cortical territories, with the risk of overlooking fine-scale heterogeneity and local variability in functional motifs. Our imaging approach bridges the gap between large-scale, low-resolution techniques and high-resolution, small-scale methods by providing both cellular resolution and millimeter-scale coverage within a single experimental framework.

An illustrative example of how this classical two-stage strategy has shaped our current understanding of functional organization in the primate brain is the combination of intrinsic imaging and classical 2p microscopy in the marmoset auditory cortex. Intrinsic imaging studies have consistently reported smooth and continuous tonotopic gradients, suggesting homogeneous frequency tuning across neighboring cortical regions^68,97^. Subsequent 2p studies similarly described local homogeneity of frequency tuning within primary auditory cortex^58^. However, these experiments did not employ soma-confined calcium indicators nor perform post hoc correction for neuropil contamination, potentially limiting their ability to resolve fine-scale heterogeneity that could be masked by background fluorescence and neuropil signals.

While our intrinsic imaging results replicated classical large-scale tonotopy maps, combining our mesoscopic cellular-resolution imaging platform with ribo-jGCaMP revealed a high local heterogeneity in neuronal frequency tuning (**Figure 3E**). Neighboring neurons frequently exhibited distinct preferred frequencies despite the presence of significant, but weak, local clustering. Importantly, the soma-confined expression of ribo-jGCaMP substantially reduced neuropil contamination and background fluorescence. This improvement also allowed us to identify neuronal populations that were either positively or negatively driven by the auditory stimuli, indicating the coexistence of mixed but distinct functional subpopulations. Together, these findings demonstrate that the smooth tonotopic gradients observed with intrinsic and conventional 2p imaging likely arise from spatial averaging over heterogeneous single-neuron tuning. By achieving reduced neuropil contamination and maintaining cellular resolution across large-scale FOVs, our approach reveals a finer-grained and more heterogeneous organization of frequency tuning that was previously obscured by methodological limitations.

In addition to areas with high local tuning heterogeneity such as the auditory cortex, the interpretation of intrinsic imaging results is also ambiguous at the boundaries or transition regions between cortical areas with different tuning properties. In these transition regions, gradual signal changes can arise from multiple factors that are not directly related to differences in selectivity, including a reduced density of responsive neurons or decreased response amplitudes of individual cells. Mesoscopic cellular-resolution imaging is therefore essential for studying how neuronal tuning and response properties change across millimeter-scale transitions between cortical areas.

By performing mesoscopic recordings across the transition zone between auditory and visual cortices, we directly captured the shift from visual to auditory tuning and identified neurons with cross-modal response profiles. Our cellular-resolution measurements revealed a complex spatial organization of auditory unimodal, visual unimodal, and multimodal neurons (**Figure 4F**). These neurons appeared as individually intermingled cells, small local clusters, and larger millimeter-scale regions dominated by a single modality. This diversity and richness of spatial tuning organization could not be resolved with intrinsic imaging alone.

Moreover, the spatial distributions of unimodal and multimodal neurons varied substantially across different parts of the transition zone and could therefore have been easily overlooked when using only a limited number of smaller FOVs or linear electrophysiological recording tracks. Although repeating recordings to cover larger cortical areas or volumes is theoretically possible, imaging the largest volume acquired here (4 × 4 × 0.3 mm³ spanning 15 planes) with a conventional 2p microscope with a single-plane FOV of 500 × 500 µm would have required 960 sequential acquisitions. Beyond the substantial experimental burden, such sequential sampling is constrained by factors including animal engagement, stimulus novelty, and the long-term stability of indicator expression and optical window clarity across recording days.

In contrast, the large spatial scale of our recordings enabled not only the capture of diverse spatial arrangements of tuned neurons, but also the continuous coverage of entire functional areas at cellular resolution. This capability allowed us, for the first time in primates, to compute large-scale, 3-D continuous cortical tuning maps (e.g., preferred direction maps and local tuning heterogeneity maps, **Figure 5F**) directly computed from individual neuronal responses.

In area MT, these cortical tuning maps revealed local clustering of direction tuning into repeated columnar motifs, or hypercolumns, with a diameter of ∼500 µm (**Figure S7Eii; Figure S8Fii**). This value is comparable to the ∼575 μm repeat distance of orientation columns in marmoset V1^98^, suggesting a similar spatial scale of functional column organization across cortical areas. We observed that the direction columns in marmoset MT are laterally tiled, arranged around pinwheel centers as previously reported in galago monkeys^76^, and across direction fractures, as previously reported in *Cebus* monkeys^73^. Similar 2-D pinwheel-like maps have previously been shown in maps of orientation tuning in V1 using low-resolution imaging approaches in other primates^5^, cats^81^, and ferrets^82^. While these approaches have provided valuable insights into average tuning properties over large cortical regions, they rely on a top-down strategy that uses coarse spatial signals, population averaging, and spatial smoothing to generate 2-D maps serving as a proxy for possible, but not certain, neuronal tuning in a region.

By contrast, our volumetric mesoscopy platform provides direct access to trial-by-trial responses of individual neurons across large cortical volumes and enables the construction of 3-D tuning maps using a bottom-up approach. These maps provide simultaneous access to summary metrics that render them interpretable across large territories while the underlying single-neuron data remain fully accessible. For instance, two regions sharing the same preferred direction can be further distinguished by a local tuning heterogeneity map that reveals whether that average preference reflects a homogeneous population or neurons with widely varying tuning (**Figure 2L,M**). More broadly, this direct access to single-cell responses allows experimenters to determine how maps should be generated to best address specific hypotheses. Importantly, unlike low-resolution methods that average across all neurons indiscriminately, our approach permits the selection of neuronal subpopulations based on their functional response properties, enabling the generation of distinct maps for different functional classes of neurons. Looking forward, future experiments could combine our imaging platform with population-specific promoters to drive the expression of genetically encoded calcium indicators in specific neuron classes, such as excitatory and inhibitory neurons, to functionally dissect their organization and tuning.

Importantly, this framework and our imaging platform also enable direct analysis of relationships between large-scale functional organization and single-neuron tuning properties, such as the negative correlation between neuronal DSI and local tuning heterogeneity. We thereby confirmed in marmoset MT a similar result reported previously in cat V1^99^, where cells in pinwheel centers were found to have a lower tuning selectivity. One methodological difference is hat this study first localized orientation pinwheels with intrinsic imaging and then targeted pinwheels with 2p imaging. In contrast, our map organization and neuronal tuning were identified using a single technique and inferred from the same dataset.

While in some cases such as mapping of single cell tuning laborious repeated sequential recordings that tile the cortical area of interest sequentially, can in principle substitute simultaneous large recording capabilities, there are a range of biological questions that critically depend on the ability to simultaneously record large scale neuronal populations. One such example addressed here is the analysis of noise correlations, which quantify whether the activity of neurons is correlated on a trial-by-trial basis and may reflect direct or indirect connectivity. This analysis requires the simultaneous acquisition of the entire neuronal population in order to decompose each response into a trial-averaged component and a trial-specific noise residual. Our volumetric LBM recordings in MT revealed that noise correlations are highest for neighboring neurons and decay rapidly and isotropically over ∼300 µm. Beyond this range, noise correlations continue to decrease monotonically with lateral distance, although they remain positive for distances larger than 3 mm (**Figure S7Evi**). These results suggest that trial-by-trial shared variability is strongly influenced by local circuit interactions, while being embedded within weaker, long-range correlations across millimeter-scale cortical distances.

Importantly, the volumetric nature of our recordings enabled us to examine how these correlation patterns vary not only laterally but also along the depth axis. Whereas noise correlations decayed isotropically over approximately 300 µm, tuning correlations remained stable across the depth axis (100–400 µm), varying primarily along the cortical surface and displaying oscillatory patterns with a spatial period of roughly 500 µm (**Figure S7Ev**). This predominance of lateral structure is consistent with the columnar organization of the cortex^35^. However, it also points to the need for future studies that extend to deeper cortical layers, where functional patterns may differ.

Cells also exhibited local spatial clustering based on their tuning class (unidirectional, bidirectional, and pandirectional) and by the anti and ortho indices we used to further describe them. However, this clustering lacked clear periodic organization at larger scales (**Figure 7D–F**), suggesting that while the cortical architecture imposes local similarity in tuning profile, it does not tile these profiles into repeated hypercolumnar motifs in the way it does for preferred direction (**Figure 2I,J,L; Figure 5Gii**).

While previous studies of MT direction tuning have necessarily focused on the more prevalent unidirectional tuning class due to the limited throughput of classical recordings, our approach allowed us to collect responses from thousands of neurons in each recording. This large throughput enabled us not only to characterize the rarer subpopulations, but to comprehensively sample the MT population. PCA of population responses revealed that the first five PCs captured the structure of direction tuning across the population, forming interpretable bases for unidirectional, pandirectional, and bidirectional cells (**Figure 6B–C**). However, neurons did not form segregated clusters in this low-dimensional space (**Figure 6D**), suggesting a functional continuum rather than discrete cell classes. This was confirmed when neurons were characterized by their anti-index and ortho-index: large numbers of cells exhibited intermediate tuning profiles, with no discrete clusters or sharp boundaries separating tuning types (**Figure 7A–B**). The ability to simultaneously record from thousands of MT neurons was essential for revealing this nuanced functional organization, as smaller-scale studies might miss the intermediate populations that bridge traditional functional categories.

The distinct stimulus-encoding strategies implied by this continuum were further revealed by examining responses to moving sinusoidal gratings. Narrow unidirectional cells preferentially responded to gratings moving in their preferred dot direction, consistent with encoding motion direction. Bidirectional cells, however, preferentially responded to gratings moving orthogonally to their preferred dot axis, suggesting that motion direction alone does not define their selectivity. One possible interpretation is that these cells perform a temporal integration of stimulus trajectory: a moving-dots stimulus, integrated over time, traces an oriented path that could be encoded as a spatial feature. Under this view, bidirectional cells may respond to the oriented spatial structure that emerges from motion over time, linking them more closely to orientation-selective mechanisms than to classical direction encoding.

Understanding how emergent properties arise in neuronal systems requires recording from sufficiently large populations while preserving single-cell resolution. Emergent structure does not manifest at the level of individual neurons or single trials alone, but through the interactions and spatial organization of many units across cortical space. At the same time, identifying these patterns demands that the heterogeneity of individual responses be retained: over-smoothing or under-sampling risks obscuring the very distinctions that give rise to population-level organization. By capturing thousands of cells simultaneously across extended cortical territories at single-cell resolution, our approach provides simultaneous access to both fine-grained cellular tuning and large-scale functional architecture within a single experimental framework. This dual-scale access reveals how computations and functional organization emerge from the collective properties of individual neurons and opens new possibilities for dissecting the principles governing cortical dynamics at unprecedented scales in the primate brain.

## METHODS

### Animal subjects and surgical procedures

All experimental procedures were approved by The Rockefeller University Institutional Animal Care and Use Committee and were performed in accordance with guidelines from the US National Institute of Health. Four male marmosets and one female marmoset aged 2.2–4.9 years were used. Surgical procedures were performed with the anesthetized animal, while its temperature, heart rate, SPO2, and CO2 were monitored. Animals were allowed to recover for at least 1 week between each of the surgical procedures. The first surgical procedure consisted of the removal of skin and muscle around the dorsal, lateral, and posterior areas of the skull, followed by the layered formation of a semi-transparent resin cement (C&B Metabond) headcap incorporating a custom stainless steel headpost. The thickness of the cement at the dorsal areas and surrounding the headpost was 5–10 mm, while a thinner 0.5–2 mm layer was applied to the lateral sides of the skull. The second surgical procedure consisted of drilling the cement, performing a craniotomy of 11 mm diameter, removing the underlying dura mater, and implanting a recording chamber with a removable imaging window similar to a previously described chamber design^50^. The third surgical procedure consisted of viral injections. Borosilicate glass capillaries (World Precision Instruments) were pulled (Sutter, P-2000) and beveled at ∼30° to obtain pipettes with 20–30 μm tips. Pipettes were backfilled with mineral oil, loaded onto injectors (World Precision Instruments, Nanoliter 2020), and front filled with 4,500 nL of the injection solution at a rate of 20–150 nL per min. The injectors were mounted on manual micromanipulators (World Precision Instruments) and magnetic stands. After removing the imaging window, two injectors were positioned orthogonally to the window and used to perform simultaneous injections. The pipettes were first positioned so their tips touched the surface of the brain, after which we advanced 500 μm, and injected 500 nL at 100 nL per min. We then advanced another 500 μm to a total depth of 1000 μm and performed a second injection with the same volume and speed. After this, we waited for 10 min before withdrawing the pipettes and moving to the next injection site. Injections were targeted to sites laterally separated by 0.5–1.0 mm and a total of 25–40 viral injections were performed in each animal.

The injected solution consisted of a mix of two viral solutions, 0.05% fast green FCF dye (Sigma-Aldrich, F7258-25G), and artificial cerebrospinal fluid (NaCl 125 mM, KCl 2.5 mM, NaHCO3 26 mM, NaH2PO4 1.25 mM, CaCl2 2 mM, MgCl2 1 mM, glucose 26 mM; described at https://doi.org/10.1101/pdb.rec092353). All injected solutions contained AAV2/9-Thy1S-tTA (Addgene 97411; produced by Vigene Biosciences) and either AAV2/9-TRE3G-jGCaMP8s (Addgene 239673), AAV2/9-TRE3-ribo-L1-jGCaMP8m (Addgene 239678), AAV2/9-TRE3-ribo-L1-jGCaMP8s (Addgene 239676), or AAV9-TRE3G-soma-jGCaMP8s (Addgene 239679). In the injecting solution, the final concentration for each virus was 0.7–1.0 × 10^13^ gc/mL. Doxycycline was periodically administered in single 0.5–20.0 mg/kg oral doses to temporarily reduce expression levels. In the process of testing different indicators and targeting strategies, we also created the following viruses for which data are not included in this study: AAV2/9-TRE3G-jGCaMP8f (Addgene 239674), AAV2/9-TRE3G-jGCaMP8m (Addgene 239675), AAV2/9-TRE3-ribo-L1-GCaMP8f (Addgene 239677), AAV9-TRE3G-soma-jGCaMP8f (Addgene 239681), and AAV9-TRE3G-soma-jGCaMP8m (Addgene 239680).

### Experimental setup

We designed and built a versatile and imaging-compatible setup where awake animals can be placed inside a marmoset holding chair, head-restrained, presented with auditory or visual stimuli, undergo eye-tracking and face-recording, and receive rewards. The setup allowed ample access to the lateral sides of the skull and can be easily modified to record from imaging windows on the occipital region. All stimulus, eye-tracking, and reward components were mounted on the same plate as the head-fixing clamp and the chair, which was mounted on two goniometers (Owis, TP 150-20-20-243), a vertical translation stage (Thorlabs, VAP4/M), and a rotation stage (Thorlabs, RP03/M). This arrangement allowed easy alignment of the imaging window with the microscope objective while maintaining the alignment of the experimental components across subjects.

For the setup, only one component was custom-made and machined (Xometry), while all other components were commercially available or 3-D printed in-house. The 3-D–printed components were designed using Autodesk Inventor and printed on an Ultimaker S3 using tough-black PLA material. The eye-tracking module consisted of a 45-degree hot mirror (Edmund Optics, 43-958) in front of the animal, two LED infrared lights (WEILAILIFE, WEI-IR-04), and a camera (Teledyne, Chameleon3), connected to a standalone computer. The position and size of the pupil were determined online using a custom-modified version of EyeLoop^100^ and output as analog channels to the stimulus computer using a DAQ (National Instruments, USB-6001). Liquid rewards (1:2 apple juice–water or 1:7 maple syrup–water) were delivered at randomly-distributed intervals from 2-8 seconds using a syringe pump (New Era Pump Systems, NE-1000).

### Stimulus presentation

Visual stimuli were presented using a display (ASUS, XG16AHPE) with a refresh rate of 144 Hz. The screen size was 34.5 cm × 19.5 cm and it was centered to the animal’s eyes and 30 cm away, covering a visual field size of 60° × 36°. For intrinsic imaging, we presented 0.4°-diameter white dots moving radially from the center of the screen over a black background^68^. During each 20 s cycle, the instantaneous speed of all dots started at 0°/s for the first 2 s, then linearly increased to 16°/s during the next 6 s, stayed at 16°/s for the next 4 s, linearly decreased to 0°/s during the next 6 s, and stayed at 0°/s for the last 2 s of the cycle. For calcium imaging, stimuli were generated using custom Python code and PsychoPy^101^. The moving-dots stimuli consisted of white and black dots moving at a constant speed over a gray background. The lifetime of the dots was 0.25 s, after which they were re-positioned to a new random location on the screen. Trial durations were 1–2.5 s and inter-trial durations were 1–1.5 s. We also presented moving sinusoidal gratings of a spatial frequency of 1 cycle per degree of visual angle and a temporal frequency of 4 cycles per second. Room lights were on during intrinsic imaging but were off during calcium imaging.

Auditory stimuli were transmitted from the stimulus computer as analog signals via a DAQ (National Instruments, PCIe-6321) to an amplifier (Tucker-Davis Technologies, SA1), which then drove two speakers (Tucker-Davis Technologies, MF1). The speakers were placed on each side of the marmoset at chest level and angled 45° relative to the ears, ensuring access for the imaging objective to the optical window. For intrinsic imaging, the audio signals were previously published^68^ and consisted of either bursts of white noise or descending sequences of pure tones. For localizing broad tonotopy in auditory cortex, we performed intrinsic imaging while presenting 20 cycles of a repeating 20 s-long trial. Each trial consisted of 2.7 s of silence, followed by a series of pure tone pips descending in frequency from 28,160 Hz to 440 Hz in one-semitone steps, and ending with 2.7 s of silence. Intrinsic responses were corrected for hemodynamic delay (3.5 s). To generate a map of tonotopic preference, we averaged the intrinsic signals to the high-frequency tones during an early time-window (+1 to +5 s after stimulus onset) and subtracted to the responses to low-frequency tones during a late time-window (−5 to −1 s before stimulus end).

For calcium imaging, the auditory white noise and tones were generated using custom Python code, were presented with and without amplitude-modulation (32 Hz, 100% modulation depth) and included a 20 ms ramp at the beginning and end of the sound. . Five marmoset call types (cry, trill, twitter, trill-phee, and phee) from three subjects recorded at postnatal days 94, 164, and 305, yielding 15 calls total, were included as auditory stimuli^102^. Signal amplitudes were calibrated using a sound level meter (General Tools, DSM403SD) so that they would have the desired intensity level. In the recordings shown in Figure 3, tones and white noise were presented with and without amplitude modulation, and all auditory stimuli were presented at 80 and 90 dB. In the recordings shown in Figure 4, tones were presented without amplitude modulation, and all auditory stimuli were presented at 90 dB.

### Intrinsic imaging

We built a cross-polarized intrinsic imaging microscope^68^ equipped with a green LED (Thorlabs, M530L4), a 2× objective (Thorlabs, TL2X-SAP; 0.1 NA, 56.3 mm WD), and a CMOS camera (FLIR, GS3-U3-23S6M-C Grasshopper 3) at 80 frames per second. The light intensity was determined by a level at which the brightest pixels (typically in the center of the window) had values of ∼90% of the saturation limit of the camera.

### Calcium imaging

We used a 2p random access mesoscope^10^ (Thorlabs, MESOSCOPE) equipped with a 2.7 mm WD objective with a 0.6 excitation NA and a 1.0 collection NA. Excitation was provided by an optical parametric chirped-pulse amplification laser (Class 5 Photonics, White Dwarf hybrid pumped by a Coherent Monaco) emitting at 960 nm and with repetition rates ranging from 3.9 to 4.9 MHz. The objective was angled 70–90° relative to the body axis of the marmoset and a 1:1 mix of centrifuged ultrasound gel and water was used as imaging media. For multi-plane Light Beads Microscopy recordings, we re-routed the excitation laser beam through a multiplexing cavity that generated 15 ‘beads’ per laser pulse. Each bead focused ∼20 µm shallower than the previous one and had 11% less power, compensating the effects of scattering and maintaining homogeneous SNR across planes. Beads arrived at the imaging sample with a temporal delay of 13.8 ns, allowing us to de-multiplex the signals. Excitation powers measured after the objective were 30–60 mW for single-plane recordings and 200–300 mW for LBM recordings. Point spread functions were collected using 0.5 µm diameter fluorescence beads and ranged 1 to 1.6 µm laterally and 8 to 13 µm axially. Lateral pixel spacing ranged from 2 to 3 µm across recordings unless otherwise stated.

### Motion correction and cell segmentation

We used the suite2p analysis package^69^ (version 0.14.4) to perform motion correction and cell segmentation. For cell segmentation, we set the parameter ‘anatomical_only’ to 3, effectively using Cellpose^103^ to perform an automated anatomical segmentation of the “enhanced mean image” (a high-pass version of the mean image). This resulted in a reliable and systematic method to obtain ROI footprints over cell soma regardless of the SNR of the recording. Since Cellpose was only used to seed and initialize ROIs and not to determine whether an ROI represented an active or responsive cell, we set Cellpose thresholds to their minimum values (cellprob_threshold = –6, flow_threshold = 0). The cell diameter in pixels was determined for each recording by dividing 13 µm by the pixel size to match the approximate median diameter of marmoset cortical neurons of 13.1 µm (ref. ^96^).

### Photon count estimation

To compare fluorescence indicators across imaging regions, ROI signals from Suite2p were converted from arbitrary units (AU) to photon units using a detector conversion factor of 126 AU/photon. The conversion factor was estimated by fitting the shot-noise model Var(B) = g · E[B] + b to the dimmest pixels identified from the temporal mean image (lowest 0.1th percentile by mean intensity), where biological fluorescence variance is minimal and the signal is most likely to be shot-noise limited. For each selected pixel, the mean and variance across all frames were used in the regression.

### Data analysis

#### Activity normalization

The spatial footprints, raw fluorescence signal (F_raw_), and neuropil signal (F_neu_) of all segmented ROIs were imported and analyzed using Python custom code. To prevent negative fluorescence values, fluorescence values of all ROIs in an imaging plane were corrected by subtracting the minimum fluorescence value across all ROIs and frames (typically −20 to −30). For each ROI, the neuropil signal was multiplied by 0.7 and subtracted from the fluorescence signal. In volumetric recordings, lateral offsets between imaging planes were determined by calculating the peak cross-correlation between adjacent planes. These offsets were used to update ROI spatial footprints and align mean images. Starting from the deepest plane and proceeding upward, ROIs were merged across adjacent planes (and for up to 4 planes) if their centers were 10 µm or less in XY coordinates, and the activity of the neurons had a pairwise Pearson correlation coefficient > 0.15. Animal movement was quantified as the sum of the absolute pixel shifts from Suite2p’s motion correction. Frames with movement exceeding 4 pixels were discarded by replacing fluorescence values with NaNs for all ROIs. We computed the fluorescence signal with neuropil subtraction for each cell as F_cell_ = F_raw_ − 0.7·F_neu_ + 0.7· F_neu_median_, where F_neu_median_ represents the median F_neu_ across all frames^88^. The baseline fluorescence of each ROI (F0) was calculated as the 40th percentile within a rolling 60-second temporal window of the frames in which no stimuli were being presented. The noise level (σ) of each ROI was computed as the robust standard deviation^104^ (1.4826 times the median absolute deviation) of the fluorescence values below F0. For each ROI, fluorescence traces were noise-normalized to compute Z activity as:

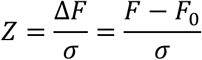

### Active-cell criteria

ROIs were classified as active if they met both of the following criteria:

1. Peaks (99th percentile) of Z activity larger than 3.
2. Peaks (99th percentile) of the dF activity larger than 3 times the recording noise, calculated as the 0.1st percentile of the dF peaks of all ROIs segmented.

We used the 99^th^ percentile rather than the absolute peak to obtain an estimate of high-amplitude activity that is less sensitive to outliers, motion artifacts, and variable frame counts across recordings and establish an exclusion criterion that is robust across recordings with different indicators, animals, frame rates, recording durations, and imaging modalities.

### Response calculation

Fluorescence activity was structured as ROIs × conditions × trials × frames. For each ROI and stimulus condition, we computed the median across trials to obtain a robust estimate of the typical response, then averaged across frames within the response window to yield a single response value per ROI per condition.

### Responsive-cell criteria

ROIs were classified as responsive if they met all the following criteria:

1. Active-cell criteria
2. ANOVA test – A one-way ANOVA across frames (grouped by stimulus condition, including blank condition) yielded p < 0.01.
3. Z activity threshold – The median response to at least one stimulus condition exceeded |Z| > 1.

### Response reliability

To assess response reliability, we followed the approach of Vinken et al.^105^.For each ROI, trials for a given stimulus condition were randomly split into two equal halves. The average response in each half was computed across trials, yielding two response vectors. Their correlation was then calculated, and this process was repeated across all possible splits to obtain the average correlation *r*. The Spearman-Brown correction was applied to estimate reliability for the full trial set:

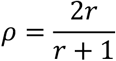

### Direction tuning

For each cell, the preferred direction *α* was calculated as:

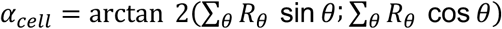

Similarly, for each cell, the preferred axis β was calculated as:

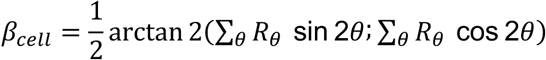

To assess direction selectivity tuning of each cell, we followed the approach of Pattadkal et al.^106^ and calculated the direction selectivity index (DSI) as:

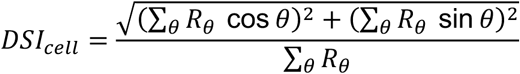

where *R_θ_* is the mean response to stimuli moving in direction *θ*.

Similarly, for each cell the axial selectivity index (ASI) was calculated as:

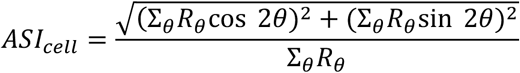

### Continuous Cortical Maps

For each recording, a 3D voxel grid spanning the cortical volume was generated with dimensions determined by the recording configuration. The grid lateral resolution was typically set to create voxels with approximately 2 µm lateral spacing. For multi-plane recordings, the z-dimension used 30 voxels (approximately 10 µm axial spacing); single-plane recordings used a single z-layer.

For each voxel, all ROIs within a sphere of influence with radius *r*= 200 µm were identified. The contribution of each ROI to a voxel’s tuning properties was weighted by distance:

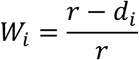

where *d_i_* is the Euclidean distance between the voxel and ROI *i*. Voxels were included only if the sum of distance weights exceeded a threshold of 5 (single-plane recordings) or 25 (LBM multi-plane recordings), ensuring reliable estimates based on both the number and proximity of contributing ROIs.

#### Direction Tuning Maps

The preferred direction of each voxel was computed as the DSI-weighted circular mean of neighboring ROIs:

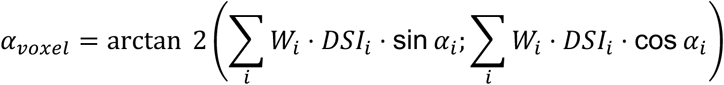

where *W_i_* is the distance weight, *DSI_i_* is the direction selectivity index, and *α_i_* is the preferred direction (0 to 2*π*) of ROI *i*.

#### Local Tuning Heterogeneity Maps

For each voxel, local tuning heterogeneity *H*was computed as:

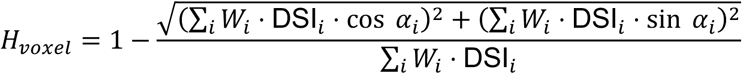

This is equivalent to 1 − *R*, where *R* is the DSI-weighted mean resultant length^107^. The numerator is the magnitude of the weighted vector sum of preferred directions, and the denominator normalizes by the total DSI-weighted contribution, so that *R*reflects directional coherence independently of the number or selectivity of contributing neurons. *H* ranges from 0 (perfect local alignment) to 1 (uniform circular distribution). Voxels exceeding the 95th of *H* were thresholded and overlaid on the direction tuning map to highlight regions of high local tuning heterogeneity.

#### Modality Preference Maps

To quantify the local preference between visual and auditory responses, each ROI was assigned a categorical modality label (’visual’, ‘both’, or ‘audio’) depending on the stimuli to which it responded with Z > 1, corresponding to numerical indices 0, 0.5, and 1 respectively. The modality preference of each voxel was then computed as the distance-weighted mean of these indices:

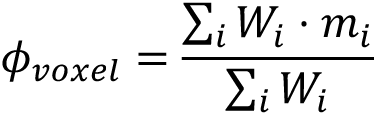

where *m_i_* is the modality index of ROI *i*(0 for visual-only, 0.5 for bimodal, and 1 for auditory-only), yielding a scalar value ranging from 0 (purely visual) to 1 (purely auditory).

### Correlation analysis

For analyzing how direction tuning and activity correlation varied with distance, we included only cells with a direction selectivity index (DSI) greater than 0.3. To examine how correlation varied with lateral distance (computed as the Euclidean distance in the x-y plane), we included only cell pairs with a depth difference of less than 50 μm. Similarly, to analyze how correlation varied with depth, we included only cell pairs with a lateral distance of less than 50 μm.

For each cell pair, we calculated their lateral and axial distances, the difference in their tuning angles, and three correlation metrics: signal correlation, trial-averaged correlation, and trial-noise correlation. Signal correlation was computed as the Pearson correlation between the pair’s normalized activity during moving-dots stimulus presentation. Trial-averaged correlation was computed as the Pearson correlation between their trial-averaged normalized activities. Trial-noise correlation was computed as the Pearson correlation between their single-trial residuals after subtracting the mean trial response.

Pairwise correlations were grouped into 50 μm distance bins. For each bin and each correlation metric, we computed the median correlation value, and standard deviation. To identify potential periodic structure in the correlation-distance relationship, we applied a fast Fourier transform to the median correlation values across binned distances. The dominant frequency component was extracted to estimate periodicity in the spatial correlation structure.

### PC subspaces for PC1–2 and PC4–5

For each neuron, we computed a combined unidirectional score as the Euclidean norm of its weights on PC1 and PC2 (sqrt(w1² + w2²)), and a combined bidirectional score as the Euclidean norm of its weights on PC4 and PC5 (sqrt(w4² + w5²)). Each neuron was then assigned to the subspace yielding its largest contribution: the unidirectional subspace (PC1–2), the pandirectional subspace (PC3), the bidirectional subspace (PC4–5), or any remaining individual PC.

## Supplemental videos index

Video S1. Single-plane 2p recording of cells expressing cyto-jGCaMP8s, related to Figure 1. Mesoscopic recording in a 2 × 2 mm FOV expressing cyto-jGCaMP8s, recorded at 3.6 Hz and played at 4× with a 4-frame rolling average. Recording is not motion corrected.

Video S2. Single-plane 2p recording of cells expressing soma-jGCaMP8s, related to Figure 1. Mesoscopic recording in a 3 × 2 mm FOV expressing soma-jGCaMP8s, recorded at 2.8 Hz and played at 4× with a 4-frame rolling average. Recording is not motion corrected.

Video S3. Single-plane 2p recording of cells expressing ribo-jGCaMP8s, related to Figure 1. Mesoscopic recording in a 2.5 × 2.5 mm FOV expressing ribo-jGCaMP8s, recorded at 2.6 Hz and played at 4× with a 4-frame rolling average. Recording is not motion corrected.

Video S4. LBM 2p recording of cells expressing soma-jGCaMP8s, related to Figure 5. Volumetric recording in a 2 × 2 × 0.3 mm FOV expressing soma-jGCaMP8s, recorded at 2.6 Hz and played at 4× with a 4-frame rolling average. Recording is not motion corrected. Planes shown in top row, from deepest to shallowest: 1, 3, 5, 7. In second row: 9, 11, 13, 15.

Video S5. Volumetric rendering of trial-averaged cell activity in response to moving dots in different directions, related to Figure 5. Recording in a 2 × 2 × 0.3 mm FOV, recorded at 2.6 Hz. The color of each cell was defined based on their preferred direction.

Video S6. Volumetric rendering of cell activity, related to Figure 5. LBM volumetric recording in a 4 × 4 × 0.3 mm FOV expressing soma-jGCaMP8s and cyto-jGCaMP8s, recorded at 1.9 Hz.

Video S7. Volumetric rendering of trial-averaged cell activity in response to moving dots in different directions, related to Figure 5. Recording in a 4 × 4 × 0.3 mm FOV, recorded at 1.9 Hz. The color of each cell was defined based on their preferred direction.

Video S8. Volumetric cortical map of direction tuning obtained from an LBM recording, related to Figure 5. Recording in a 4 × 4 × 0.3 mm FOV, recorded at 1.9 Hz.

**Figure S1.**
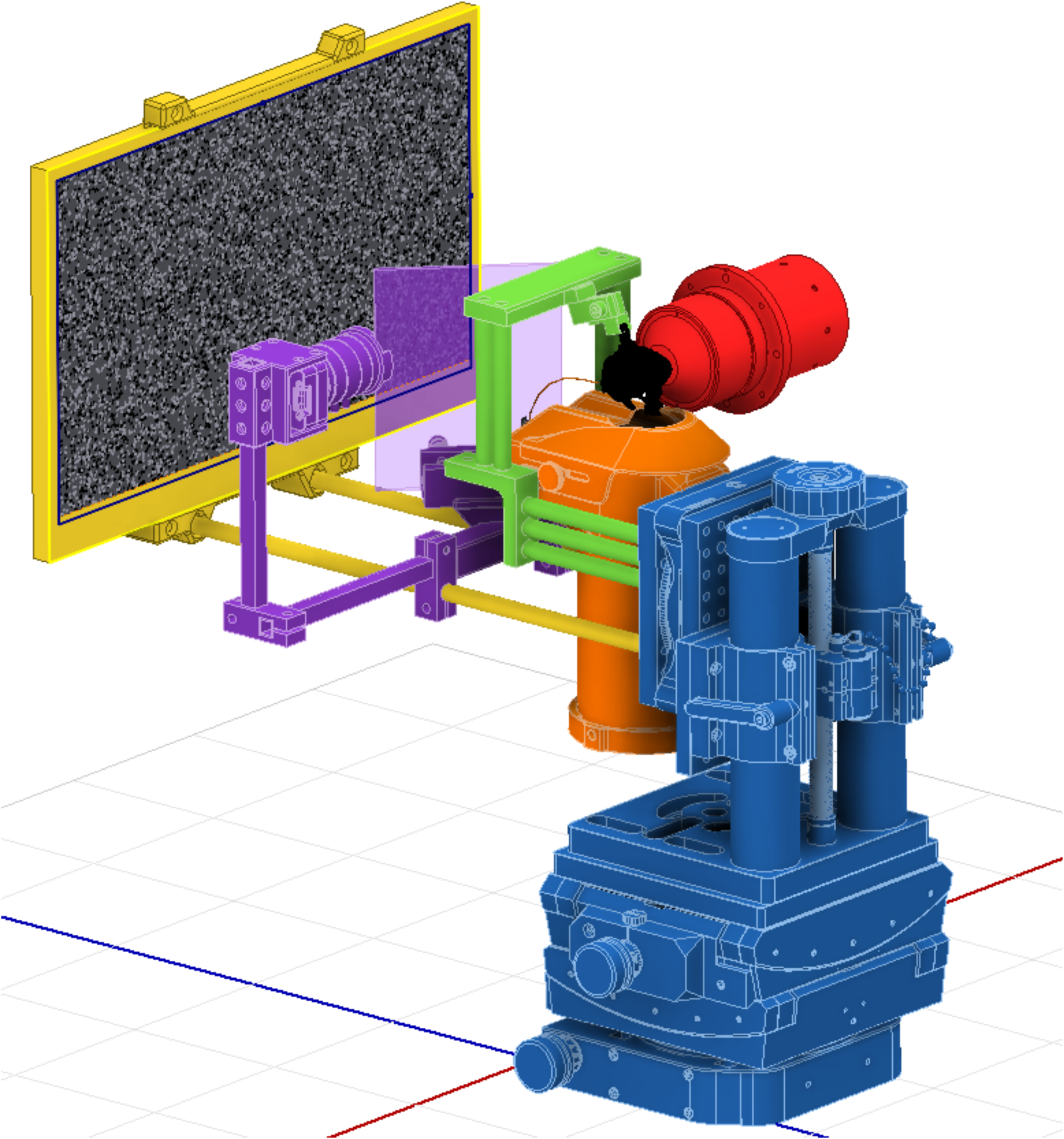
Custom marmoset imaging setup. A custom-built setup enables versatile positioning of the animal while maintaining alignment between visual stimulation and eye-tracking components. The mounting base (blue) integrates, from bottom to top, a horizontal rotational stage, goniometers, a vertical translation stage, and a vertical rotational stage. A 3D-printed marmoset chair (orange) and head-fixation system (green) are anchored to this base to secure the animal (black). The visual stimulus display (yellow) and eye-tracking camera (purple) are mounted on the same base, preserving their alignment across positional adjustments. The neckplate design of the chair provides unobstructed access for large imaging objectives (red).

**Figure S2.**
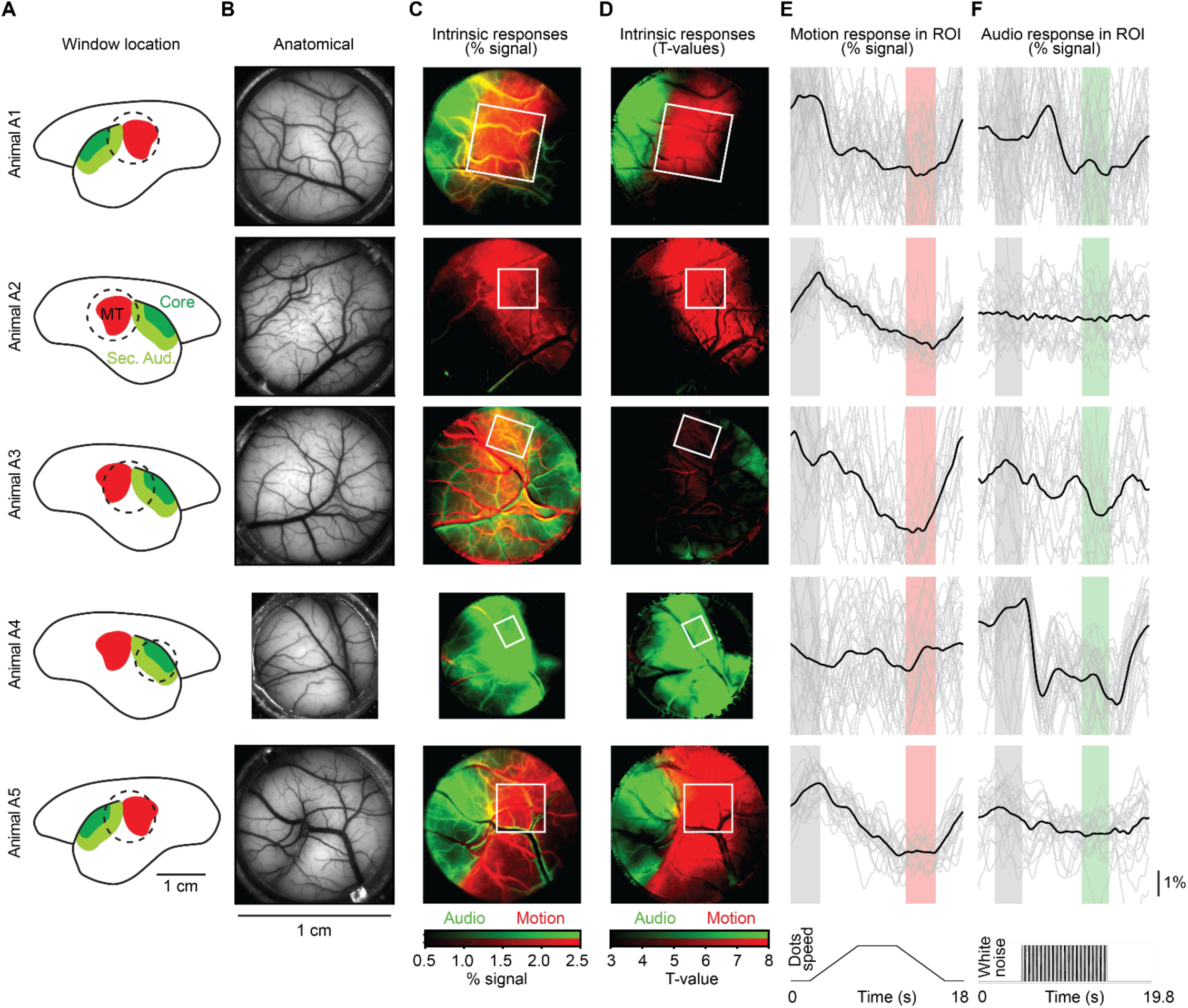
Localization and functional mapping of imaging windows using intrinsic imaging. The MT complex and auditory cortex were localized within the imaging windows based on hemodynamic responses to moving-dot visual stimuli and white-noise bursts. A. Schematic of estimated imaging window positions relative to the MT complex (red), auditory core (dark green), and secondary auditory cortices (light green), based on stereotaxic coordinates and the functional data in panels C–D. B. Raw anatomical images of the imaging windows acquired with the intrinsic imaging microscope. C–D. Functional intrinsic-signal maps showing hemodynamic responses to moving dots (red) and white noise (green), expressed as percentage signal change (C) and T-values (D). White rectangles indicate the 2p imaging FOV locations. E–F. Time courses of intrinsic responses to moving dots (E) and white noise (F) within the regions marked by white rectangles in C–D. Gray shading indicates the baseline period; red and green shading denote the response windows for motion and auditory stimuli, respectively.

**Figure S3.**
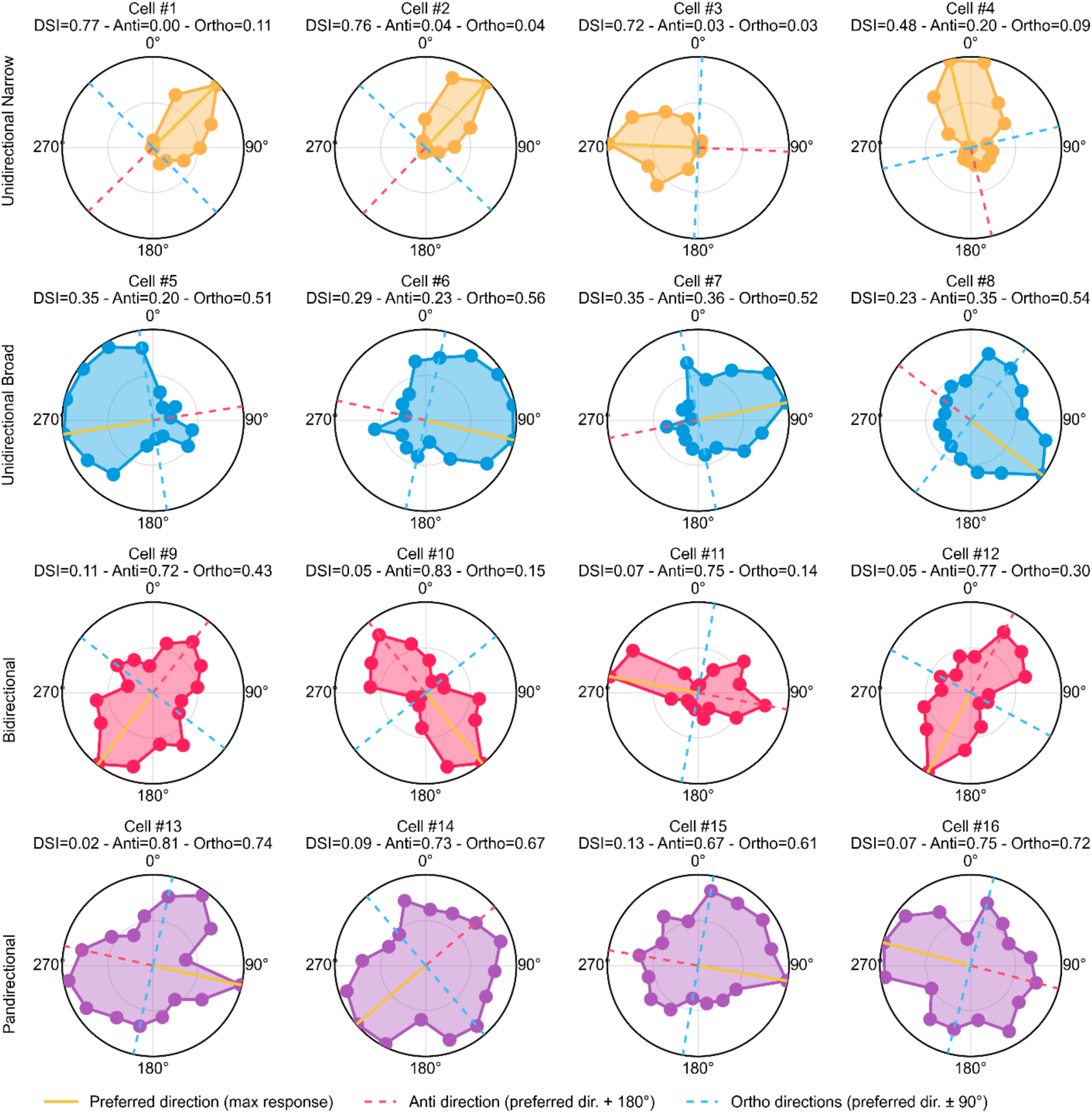
Example single-cell responses to moving-dots stimuli in MT, related to Figure 2. Polar plots of trial-averaged responses to moving dots in 16 directions for 16 example cells, arranged in four rows of four cells each. Rows correspond to the four tuning types defined in the main text, color-coded by tuning type: narrow unidirectional (top row, yellow; low anti index and low ortho index), broad unidirectional (second row, blue; low anti index, high ortho index), bidirectional (third row, red; high anti index, low ortho index), and pandirectional (bottom row, purple; high anti index and high ortho index). Within each polar plot, the orange line indicates the maximally responsive, the red dashed line the anti-preferred direction (preferred + 180°), and the blue dashed lines the orthogonal directions (preferred ± 90°). The DSI, anti index, and ortho index are reported above each cell.

**Figure S4.**
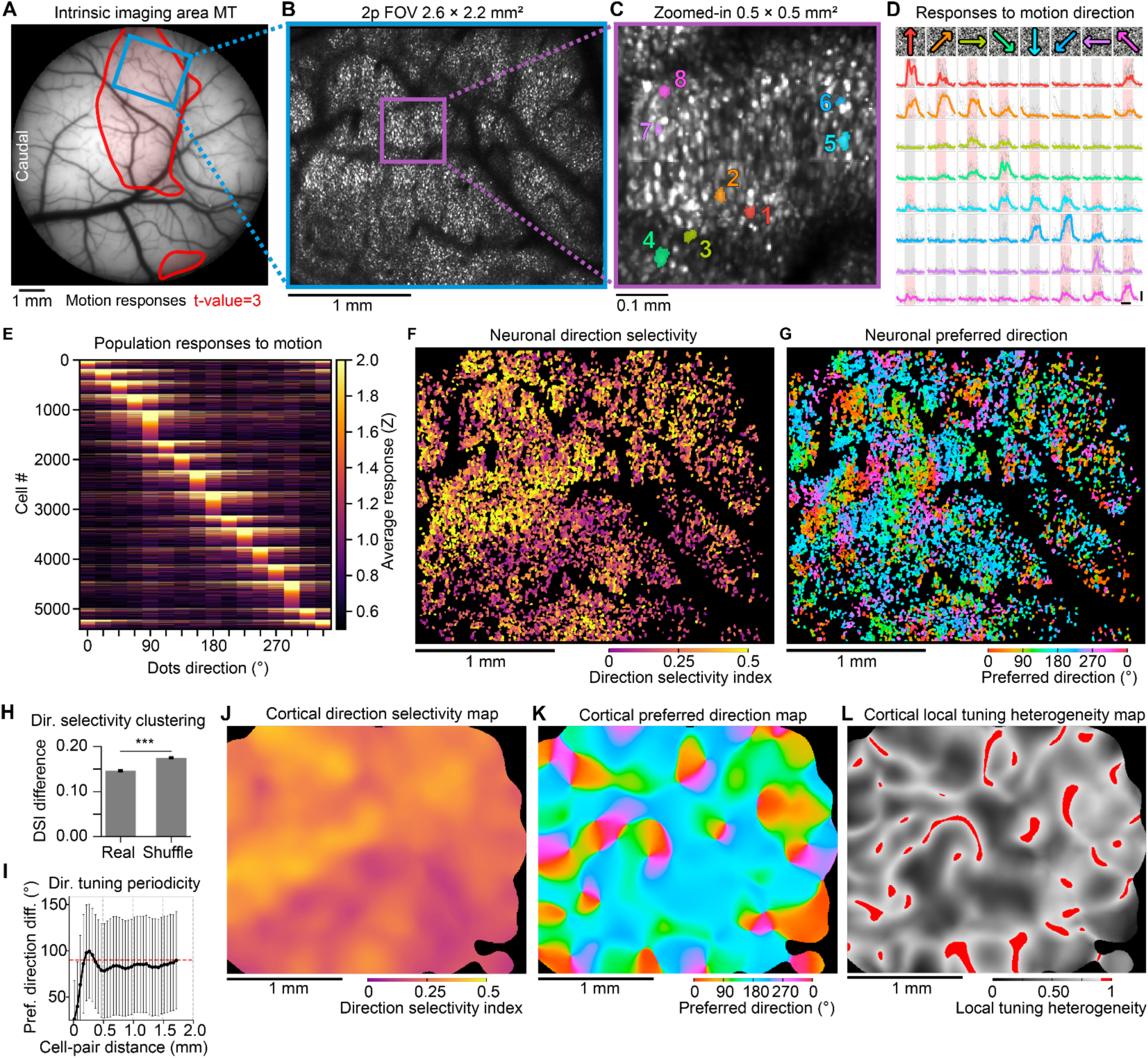
Cellular-resolution mesoscopic imaging and direction tuning in marmoset MT in a second animal, related to Figure 2. A. Functional intrinsic-signal map of an optical window showing the location of MT based on the hemodynamic responses to a moving-dots stimulus. The blue rectangle marks the mesoscopic 2p FOV shown in panel B. B. Mean image of a mesoscopic 2p recording showing dense expression of ribo-jGCaMP8s throughout the entire FOV of 2.6 × 2.2 mm² at 5.38 Hz. The purple rectangle marks the zoomed-in region shown in panel C. C. Close-up view of a 0.5 × 0.5 mm² region from panel B shows dense labeling and soma-confined expression of ribo-jGCaMP8s. The footprints of segmented cells are numbered and colored based on their preferred direction. D. Stimulus-aligned activity for the cells shown in panel C displays stimulus-locked responses to different subsets of moving-dots directions. Each row corresponds to one cell and each column to a moving-dots direction. Gray lines represent the responses to individual trials and colored lines the median response across trials, where hue corresponds to the preferred direction of the cell. Stimulus presentation periods are shaded red for responses > 1 Z or gray otherwise. Horizontal scale bar: 2 s; vertical scale bar: 5 Z. E. Heatmap of trial-averaged activity from all stimulus-responsive cells (n = 5,422 of 9,228 active cells, 58.8%), showing responses to motion stimuli. Trial-to-trial reliability was 0.70 ± 0.28 (median ± SD). Responsive cells were classified as narrow unidirectional (n = 3,789, 69.9%), broad unidirectional (n = 706, 13.0%), bidirectional (n = 591, 10.9%), or pandirectional (n = 336, 6.2%). F. Footprints of cells colored by DSI. The mean DSI across responsive cells was 0.27 ± 0.16 (mean ± SD). G. Footprints of cells colored by preferred direction, showing local clustering of directional tuning. H. Local clustering of direction selectivity, quantified as the mean DSI difference between each cell and its five nearest neighbors (0.15), was lower than expected by chance (shuffle: 0.17; permutation test, 100 iterations). I. Preferred direction differences for pairs of neurons sorted by cell-pair distance. Pairs less than 50 µm apart differed in preferred direction by 26.0° ± 41.7° (median ± SD), while pairs 200–250 µm apart differed by 97.7° ± 51.9°. The oscillatory modulation of preferred direction differences had a dominant spatial period of 437.5 µm. The red dashed line at 90° represents the expected random distribution. J. Continuous map of direction selectivity, where each pixel’s color reflects the weighted average DSI of cells within a 200 µm radius. K. Continuous map of preferred direction, where each pixel’s color reflects the weighted average preferred direction of cells within a 200 µm radius. L. Continuous map of local tuning heterogeneity, where each pixel’s brightness reflects the weighted circular variance of preferred directions of cells within a 200 µm radius. Regions with values > 0.9 are highlighted in red, marking putative pinwheel centers and direction fractures.

**Figure S5.**
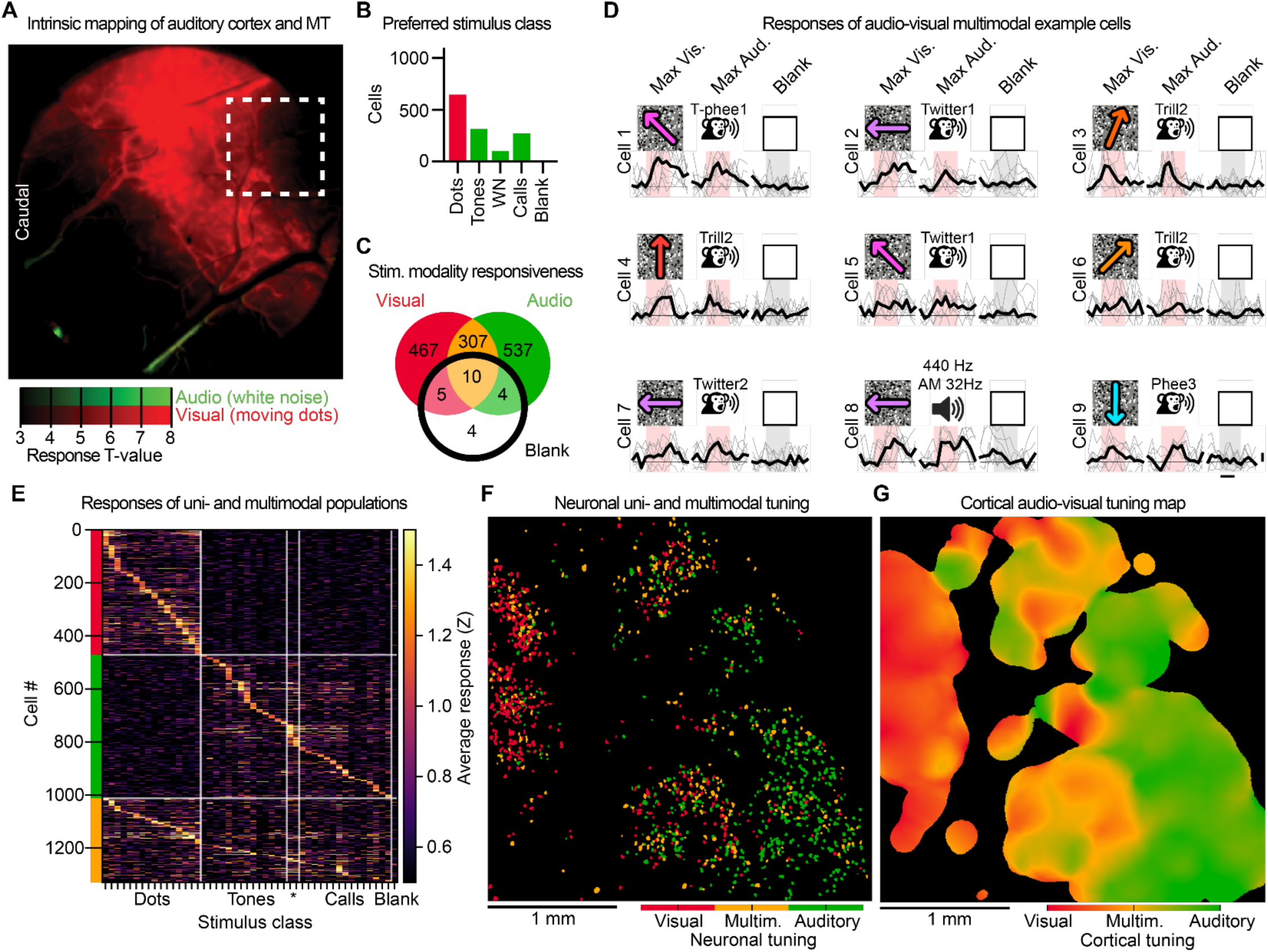
Cellular resolution mesoscopic imaging of the auditory-visual cortical border and multimodal tuning in a second animal, related to Figure 4. A. Functional intrinsic-signal map of an optical window showing adjacent regions with hemodynamic responses to moving dots (red) and white noise bursts (green). The dashed rectangle marks the 2p recording FOV from which the data in panels B–G were obtained. Scale bar: 1 mm. B. Number of stimulus-responsive cells (total n = 1,335) that responded maximally to each stimulus class: moving dots in 16 directions, pure tones at seven frequencies (440 to 28,160 Hz), white noise (WN), marmoset vocalizations (Calls), and blank. C. Venn diagram showing the number of cells that responded significantly (mean Z > 1) to at least one visual stimulus only (red, n = 472), at least one auditory stimulus only (green, n = 541), stimuli of both modalities (yellow, n = 318), or blank (black circle). D. Stimulus-aligned activity for nine multimodal example cells, each shown for three conditions: their maximum visual stimulus (left), maximum auditory stimulus (center), and blank (right). Each column corresponds to one stimulus condition. Gray lines represent individual trials and the black line the median response across trials. Stimulus periods are shaded red for responses > 1 Z or gray otherwise. Auditory stimuli are labeled by call type and caller identity, with five call types each from three different animals. Horizontal scale bar: 1 s; vertical scale bar: 1 Z. E. Heatmap of trial-averaged population responses to moving dots, pure tones, white noise (asterisk), marmoset vocalizations (Calls), and blank. Cells are grouped and sorted by modality preference: visual unimodal (red bar), auditory unimodal (green bar), and multimodal (yellow bar). F. Spatial distribution of cells colored by their modality classification: visual unimodal (red), multimodal (yellow), and auditory unimodal (green). G. Continuous map of audio-visual tuning, where each pixel’s color reflects the weighted average audio-visual tuning index of cells within a 200 µm radius, ranging from visual (red) to multimodal (orange/yellow) to auditory (green).

**Figure S6.**
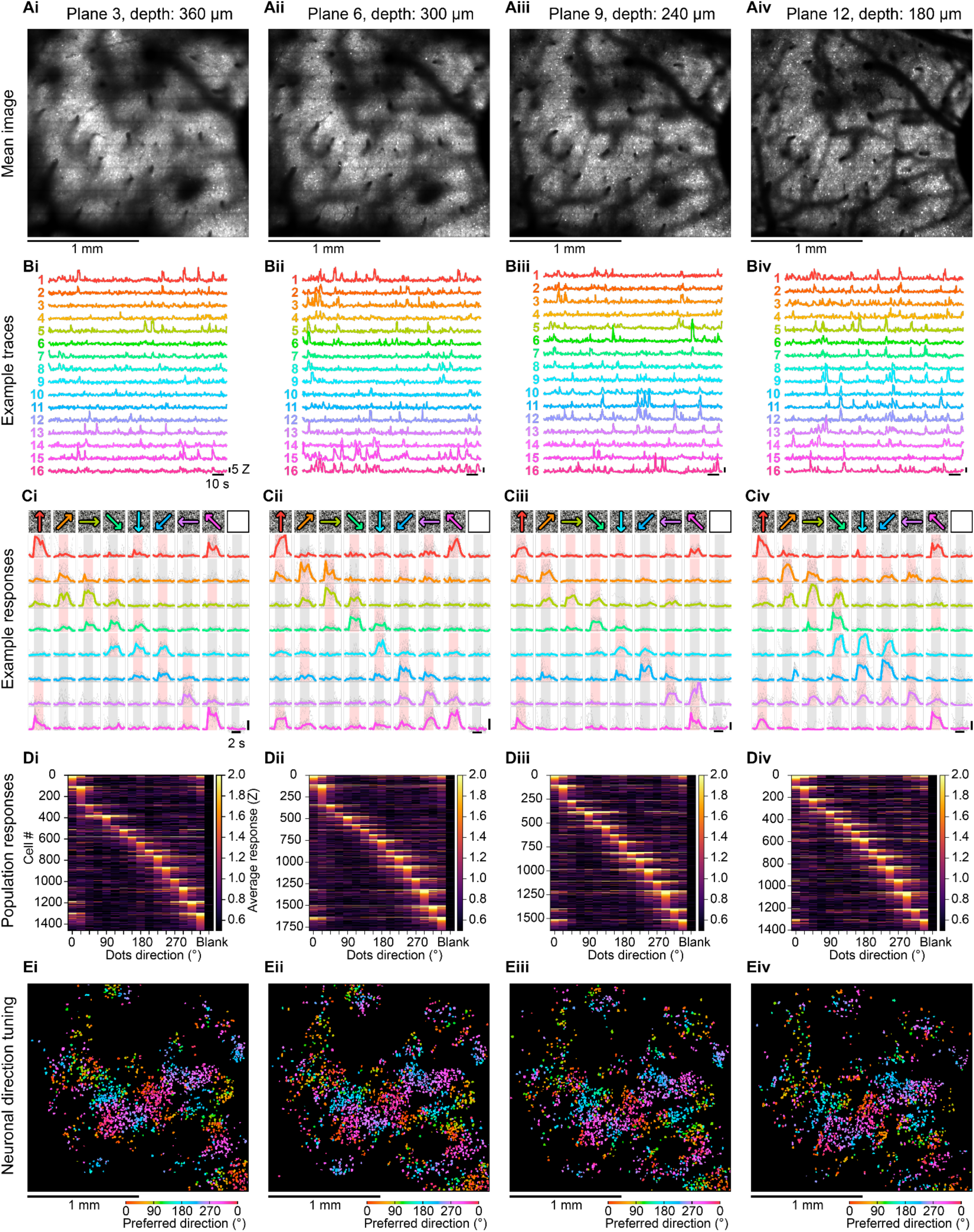
Direction tuning across LBM planes in 2 × 2 × 0.3 mm^3^ volume acquired from MT, related to Figure 5. A. Mean images of four evenly spaced planes at different depths in the LBM recording volume. B. Calcium activity traces of 16 example cells from each plane, numbered and colored by preferred direction (hue-to-direction reference as in E). Horizontal scale bar: 10 s; vertical scale bar: 5 Z. C. Stimulus-aligned activity for the 8 odd-numbered cells shown in B (cells 1, 3, 5, 7, 9, 11, 13, 15), displaying stimulus-locked responses to different subsets of moving-dots directions. Each row corresponds to one cell and each column to a stimulus condition (moving-dots direction or blank, rightmost). Gray lines represent the responses to individual trials, and colored lines represent the median response across trials where the hue corresponds to the preferred direction of the cell. Stimulus presentation periods are shaded red for responses > 1 Z or gray otherwise. Horizontal scale bar: 2 s; vertical scale bar: 5 Z. D. Heatmap of trial-averaged activity from all stimulus-responsive cells in each plane, showing robust responses to motion stimuli but not to the blank condition (rightmost column). E. Footprints of cells in each plane colored by preferred direction, showing local clustering of directional tuning consistent across depths.

**Figure S7.**
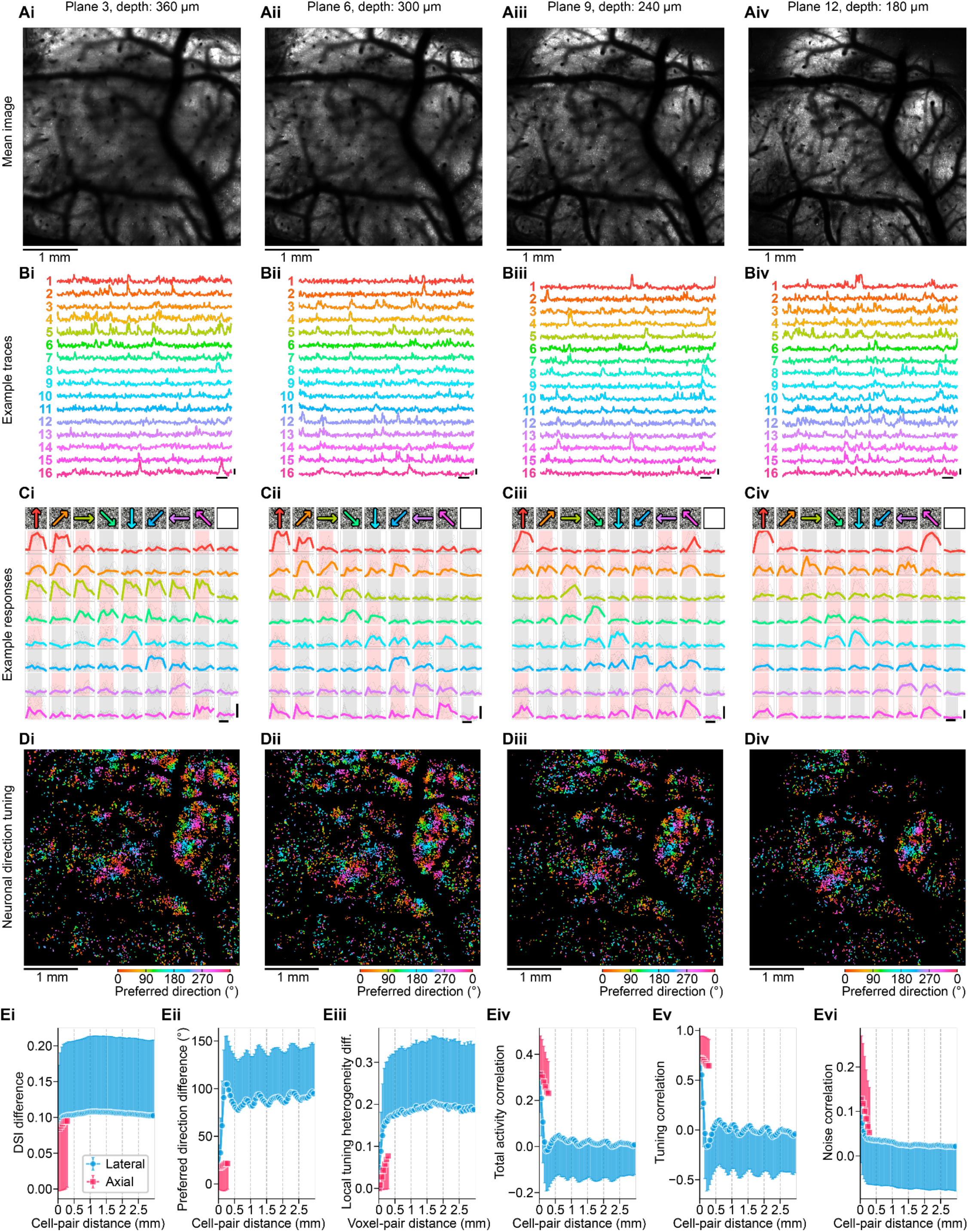
Volumetric functional characterization of direction tuning in MT using LBM in a 4 × 4 × 0.3 mm³ FOV, related to Figure 5 and Figure 6. A. Mean images of four evenly spaced planes at different depths in the LBM recording volume. B. Calcium activity traces of 16 example cells from each plane, numbered and colored by preferred direction (hue-to-direction reference as in D). Horizontal scale bar: 10 s; vertical scale bar: 5 Z. C. Stimulus-aligned activity for the 8 odd-numbered cells shown in B (cells 1, 3, 5, 7, 9, 11, 13, 15), displaying stimulus-locked responses to different subsets of moving-dots directions. Each row corresponds to one cell and each column to a stimulus condition (moving-dots direction or blank, rightmost). Gray lines represent the responses to individual trials, and colored lines represent the median response across trials where the hue corresponds to the preferred direction of the cell. Stimulus presentation periods are shaded red for responses > 1 Z or gray otherwise. Horizontal scale bar: 2 s; vertical scale bar: 5 Z. D. Footprints of all direction-responsive cells in each plane (n = 4,626, 4,995, 3,907, and 2,909, from superficial to deep) colored by preferred direction, showing local clustering of directional tuning consistent across depths. E. Pairwise-distance analysis across the volume (median ± SD) for: (i) neuronal DSI difference, (ii) neuronal preferred-direction difference, (iii) cortical local tuning heterogeneity difference across voxels, (iv) neuronal total activity correlation, (v) tuning correlation, and (vi) noise correlation. Data corresponding to lateral (XY, blue) and axial (Z, red) separations are shown separately. The modulation of preferred direction differences, total activity correlations, and tuning correlations exhibited across lateral separation had a dominant spatial period of 500 µm.

**Figure S8.**
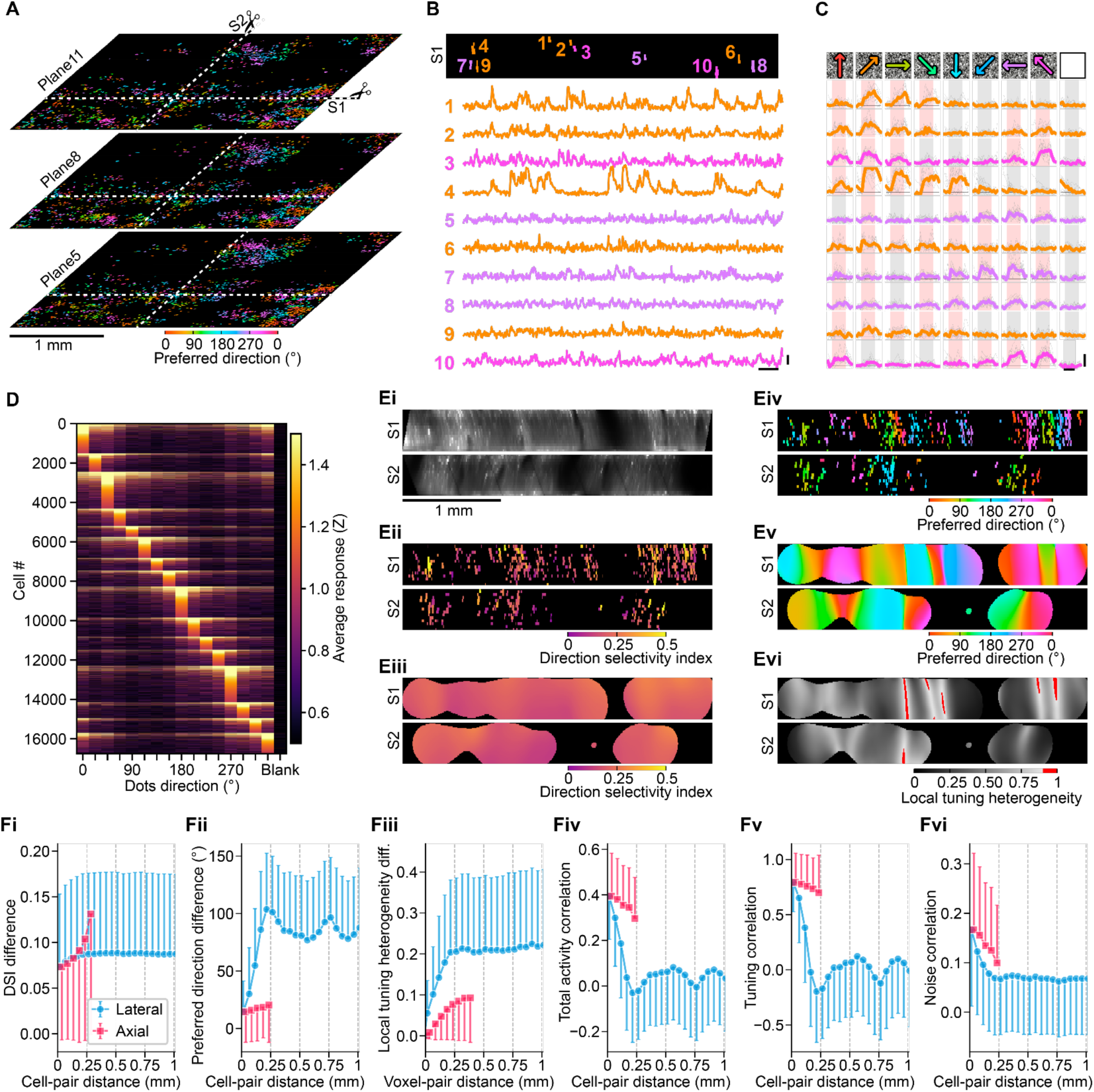
Volumetric functional characterization of direction tuning in MT using LBM in a second animal in a 3.1 × 3.1 × 0.3 mm³ FOV, related to Figure 5. A. Three of the 15 planes imaged with LBM are shown, with cells colored by preferred direction. Dashed lines indicate the Y position for the XZ slice (S1) and the X position for the YZ slice (S2) shown in S8B and S8E. B. The top panel shows an orthogonal XZ view (S1) from the recording volume where 10 footprints of selected example cells were colored by their preferred direction (color conventions as in A). The bottom panel shows the activity traces of the example neurons. Horizontal scale bar: 10 s; vertical scale bar: 5 Z. C. Stimulus-aligned activity for the 10 cells shown in panel B displays stimulus-locked responses to different subsets of moving-dots directions. Each row corresponds to one cell following the same order as in B, and each column to a stimulus condition (moving-dots direction or blank, rightmost). Gray lines represent the responses to individual trials, and colored lines represent the median response across trials where the hue corresponds to the preferred direction of the cell. Stimulus presentation periods are shaded red for responses > 1 Z or gray otherwise. Horizontal scale bar: 2 s; vertical scale bar: 5 Z. D. Heatmap of trial-averaged activity from all stimulus-responsive cells (n = 16,754 of 82,558 active cells, 20.3%), showing strong responses to motion stimuli but not to the blank condition (rightmost column). Trial-to-trial reliability was 0.61 ± 0.20 (median ± SD). The mean DSI across responsive cells was 0.20 ± 0.11 (median ± SD). E. Orthogonal views of the slices S1 and S2 showing: (i) mean image; (ii) neuronal DSI; (iii) cortical DSI map; (iv) neuronal preferred direction; (v) Continuous map of preferred direction; and (vi) local tuning heterogeneity map. F. Pairwise-distance analysis across the volume (median ± SD) for: (i) neuronal DSI difference, (ii) neuronal preferred-direction difference, (iii) cortical local tuning heterogeneity difference across voxels, (iv) neuronal total activity correlation, (v) tuning correlation, and (vi) noise correlation. Data corresponding to lateral (XY, blue) and axial (Z, red) separations are shown separately. The oscillatory modulation of preferred direction differences, total activity correlations, and tuning correlations with lateral separation had a dominant spatial period of ∼408 µm. For each datapoint, the SD is shown only in one direction to aid visualization.

**Figure S9.**
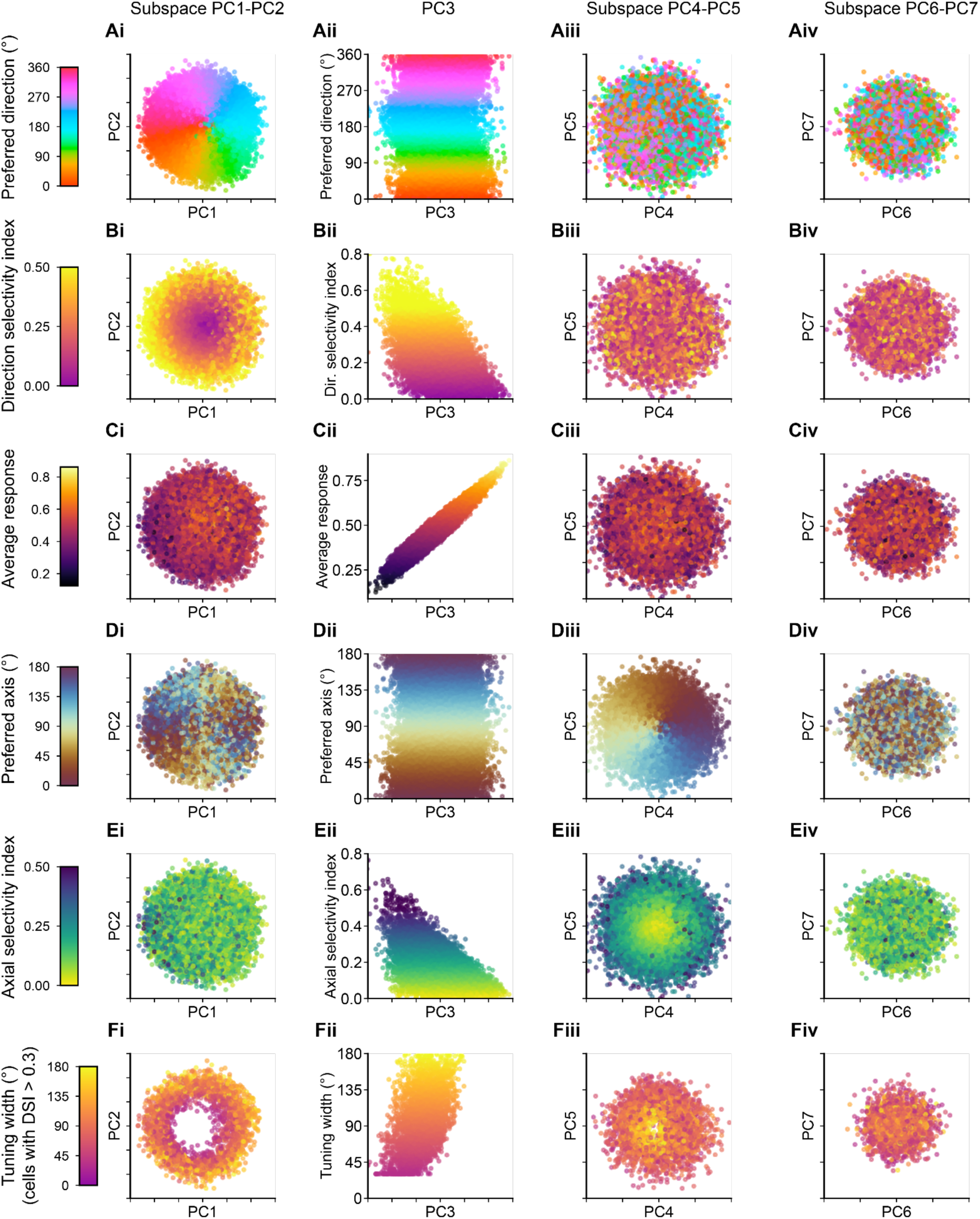
Representation of direction tuning metrics across PC subspaces, related to Figure 6. Each panel shows individual cells (dots) projected onto a PCA subspace, color-coded by a tuning metric. Rows correspond to metrics (A–F, top to bottom); columns correspond to PCA subspaces (i–iv, left to right): PC1–PC2 (i), PC3 (ii), PC4–PC5 (iii), and PC6–PC7 (iv). For 2-D subspaces (columns i, iii, iv), axes represent the two PC scores; for the single-component column (ii), the x-axis is the PC3 score and the y-axis is the metric value. Colorbars at the far left of each row define the color-to-value mapping for all panels in that row. Rows show: (A) preferred direction, (B) DSI, (C) average response magnitude, (D) preferred axis, (E) ASI, and (F) tuning width, estimated as the FWHM of a von Mises fit to each cell’s direction tuning curve, shown only for cells with DSI > 0.3. Panels Ai, Bi, Cii, Diii, and Eiii are reproduced in Figure 6.

**Figure S10.**
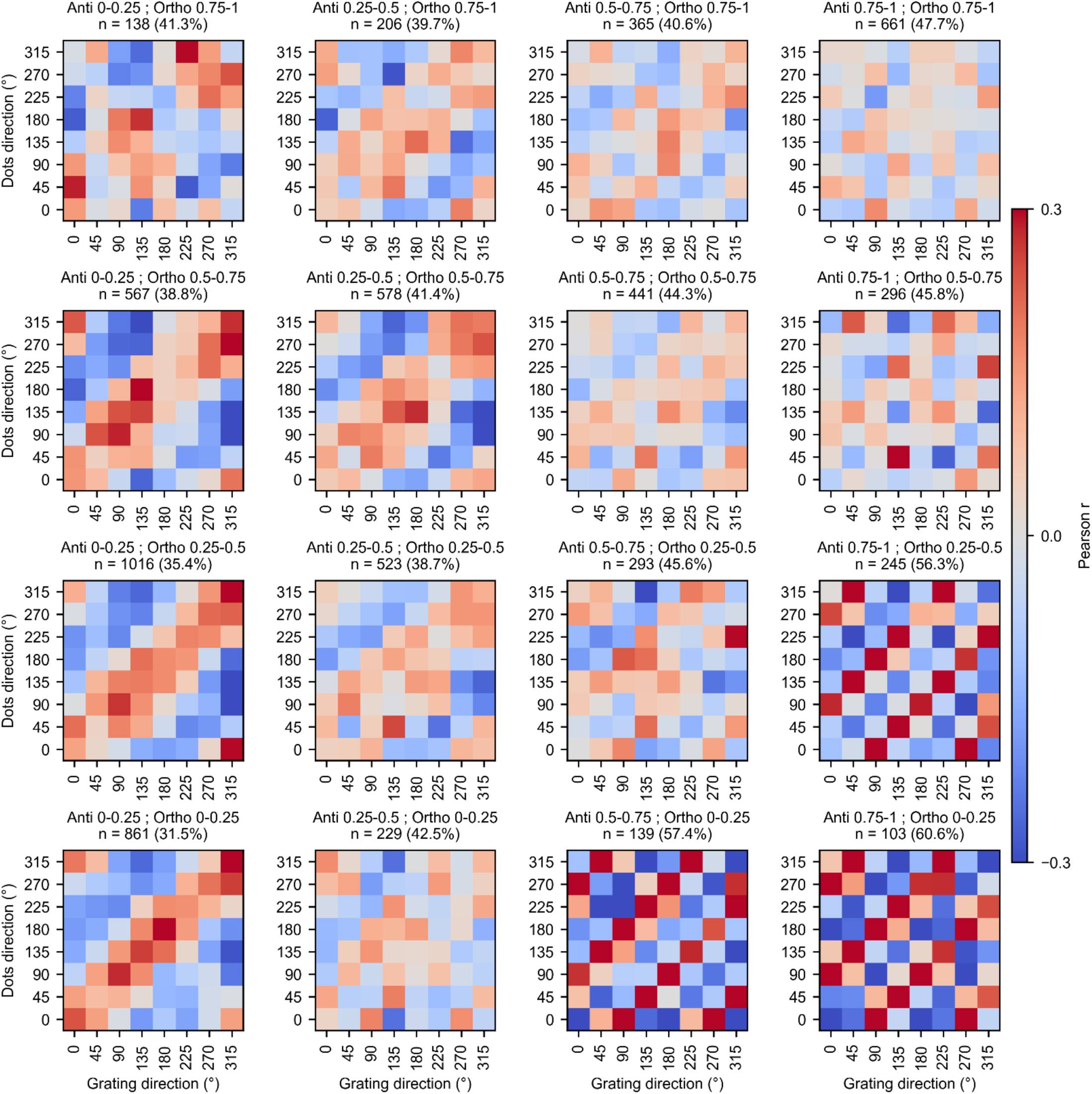
Cross-stimulus direction tuning correlations across anti index / ortho index bins. Each panel shows an 8×8 direction-by-direction Pearson correlation matrix comparing responses to moving-dots stimuli (y-axis) and moving sinusoidal gratings (x-axis), computed across cells within a given anti index and ortho index bin. The anti index and ortho index of cells were calculated based on their responses to moving-dots stimuli. For the correlation analysis, tuning vectors were zero-centered (i.e., mean-subtracted) per cell prior to correlation. Bins are arranged in a 4×4 grid with anti index increasing left to right and ortho index increasing from bottom to top. Bin titles indicate bin bounds. Cell counts and percentages indicate the number of cells in each bin responsive to both stimulus classes (Z > 1.5 for at least one direction of each), as a fraction of all dots-responsive cells in that bin.

## ACKNOWLEDGEMENTS

We thank Peer Strogies and the Precision Instrumentation Technologies (PIT) team at Rockefeller University for fabrication of mechanical components, A. Gonzalez and the Comparative Bioscience Center of the Rockefeller University for veterinary services and providing surgical assistance. Research reported in this publication was supported by the National Institute of Neurological Disorders and Stroke of the National Institutes of Health under award numbers 5U01NS126057 (A.V.), 5U01NS103488 (A.V., W.A.F.), 5U01NS094263 (A.V.), the National Eye Institute of the National Institutes of Health under award number R21EY037066 (W.A.F.) and R21EY031486 (W.A.F.); the National Institute of Mental Health of the National Institutes of Health under award number R21MH125188 (W.A.F.); the Simons Foundation under grant number 365002 (W.A.F.); the Kenneth C. Griffin Fund (A.V., W.A.F.); the Kavli Foundation through the Kavli Neural System Institute (A.V., S.W., J.D., S.O.-C., D.G.C.H.); a NARSAD Young Investigator Grant from the Brain & Behavior Research Foundation under grant number 29422 (D.G.C.H.); and funds from Bristol Myers Squibb Postdoctoral Fellowship at The Rockefeller University (J.D.), Leon Levy Fellowships at The Rockefeller University (S.W., D.G.C.H.), and Price Family Center for the Social Brain Fellowship at The Rockefeller University (S.O.-C.). The content is solely the responsibility of the authors and does not necessarily represent the official views of the National Institutes of Health.

## AUTHOR CONTRIBUTIONS

S.O.C., D.G.C.H. contributed to the conceptualization and design of the experiments. S.O.C. and D.G.C.H. conducted surgeries, performed experiments, and performed initial data analysis. B.C. assisted with surgeries and animal handling, S.O.C. performed extended analysis and generated the visualizations. T.N. supported the hardware control aspect of the imaging system and provided guidance on data analysis and software related issues. A.V. designed imaging system, J.D., F.T. and S.W. contributed to realization of the imaging system and S.O.C. to its characterization and validation. D.G.C.H., S.G., K.K.L., G.H.V., and M.F. designed and produced the viral vectors. D.G.C.H., S.W., B.C., J.D. performed experiments on an earlier version of the reported imaging platform. S.O.C., D.G.C.H., W.A.F., and A.V. wrote the paper with input from all authors. W.A.F. and A.V. conceived and led the project, conceptualized, designed and guided experiments and data analysis approach.

## Notes

### Competing Interest Statement

The authors have declared no competing interest.

